# Elucidation of the polysaccharide cryoprotection mechanism: kinetic inhibition, thermodynamic Gibbs-Thomson modulation and volumetric confinement

**DOI:** 10.1101/2025.08.27.672675

**Authors:** B. M. Guerreiro, J. C. Lima, J. C. Silva, F. Freitas

## Abstract

The emergence of gel-forming, ice-binding polysaccharides as potential candidates in cryobiology and the discovery of new structure–function relationships has fueled a knowledge convergence effort. Several polysaccharides have shown strong biological post-thaw benefits in cell cryopreservation despite some expressing contradictory ice growth anticipation, the main source of cryoinjury. The bio-based fucose-rich polysaccharide FucoPol, a current model under our scope of expertise, has further demonstrated crystal size reduction, thermal hysteresis, nucleation anticipation, nucleation stochastic narrowing and Gibbs–Thomson growth modulation effects, the latter similar to a type I antifreeze protein, in different thermodynamic settings: bulk vs. directional freezing; isobaric vs. isochoric systems; and sterile vs. biological media. Here, we undergo a critical reiteration of the consortium of findings over the years on cryoprotective polysaccharide research, rationalize several heuristic models to explain a dual nucleation behavior scenario and put forth a unifying theory to explain in which subset of conditions optimal cryoprotection may emerge from bio-based polysaccharides. We argue that gel-forming, ice-binding polysaccharides that show cryoprotective traits by anticipating nucleation and ice growth act by a combination of kinetic hampering of molecular diffusion (concentration effect); Gibbs–Thomson specific ice binding (templating effect) that induces growth modulation and size reduction; and indulge in the formation of a gel architecture of defined porosity (mesh size effect), the main initiator of a predominant pro-nucleation setting, increased stochastic determinism and the annihilation of large *r*\* nuclei that elicits a small-nuclei survivorship bias. Classical Nucleation Theory formalisms support this hypothesis under the circumstance that a change in system state, initiated by a sol–gel transition near hypothermia, must exist to drive a meaningful shift in system energetics that explains the ANTI/PRO nucleation duality observed. The molecular fractioning of kinetically hindered bulk water into fractionally partitioned gel pores reduces system scale by a mesh size constant and enables the Gibbs–Thomson ice-binding affinity condition to be satisfied when *r* = *r*_GT_. A decreased nucleation energy barrier and maximal *r*\* constraint thus invokes a predominant pro-nucleating system by enhancing nucleation susceptibility.

## Introduction

An agglomeration of research efforts on cryoprotective polysaccharides has unveiled several structure-function relationships that have enabled a better understanding of the underlying mechanisms of such molecules in sub-zero temperature biological systems. A recent meta-analysis review [1] and a polysaccharide screening [2] highlighted the importance of polyanionicity towards achieving binding interactions at the polysaccharide-ice interface to exert disruptive ice growth inhibition. Although this property is conferred not only by an elevated content of negatively charged uronic acids (UA) but also structural dispositions that maximize those interactions, mainly branched structures. Candidate polysaccharides must also adhere to solubility, biocompatibility and solution conductivity constraints, which relates to net negative charge, in order to be adequate cryoprotectants in a biological setting without inducing toxicity scenarios [3]. After a screening of cryoprotective candidates [2], those who ensured the highest biological viability were able to generate high viscosity microenvironments from non-covalent entanglement, formed a gel-state near subzero temperatures, induced an anticipation of solution freezing temperature (attenuated supercooling) and possessed ordered structures in helix or sheet forms [4]. FucoPol, a bio-based fucose-rich polysaccharide produced by *Enterobacter* A47 (Figure *SI*.*3*), which has been the model polysaccharide on our extensive cryobiological research [5], was the best performing carbohydrate polymer [6]. All these properties hinted at a compound structural contribution towards the expression of an antifreeze effect, which was most likely related to their monomeric composition. As suggested by the monomer fingerprinting in [1], cold-adapted psychrophilic polysaccharides often express above-average amounts of UA in their composition, and sometimes increased contents in neutral fucose and rhamnose. These trends were also observed for salt-adapted halophiles. Later, the synthesis of tosylated FucoPol variants that targeted the complete removal of UA from the structure showed that, although biocompatibility was preserved, it resulted in complete loss of cryoprotective potential and the inability in forming a gel-state near hypothermia due to the loss of critical non-covalent interactions by increased hydrophobicity [3]. In addition, a strong relationship between fucose content and cryoprotective function was uncovered [2]. The surmounting evidence from the cryobiological screening pinpointed UA-rich, fucose-rich polysaccharides to consistently dominate the top-tier cryoprotective performance amongst a consortium of 26 polysaccharides. However, fucose-rich polysaccharides could outperform the optimized, commercially available, stem-cell cryopreservation formula CryoStor CS5. Principal Component Analysis revealed that fucose and UA contribute towards cryoprotective outcome by different mechanisms [2], which agrees with their distinct neutral and anionic nature, respectively, but nevertheless urged a paradigm shift. The role of polyanionicity at the polysaccharide-ice interface was validated, but the presence of fucose highlighted the impact of a necessary hydrophilic-hydrophobic net charge balance. This has also been observed in natural antifreeze proteins [7–9], synthetic polyampholytes [10–13] and other glycopolymers [14, 15]. The known glycobiological role of fucose in membrane interactions [16–19], its overexpression in halophiles which require high salinity adaptation by osmoregulation [1] and its propensity to be present in top-tier cryoprotective structures [2] all pointed to fucose playing a critical role towards cryoprotective outcome, whether at the polysaccharide-membrane interface by direct membrane stabilization effects, or by indirectly driving a beneficial conformational folding arising from its inclusion. At the polysaccharide-ice interface, interfacial effects were explored by directional freezing and demonstrated that FucoPol indulges in dynamic ice shaping of growing ice dendrites, vastly decreasing their thickness and sharpness, while the highly viscous environment leads to dendritic alignment and better compactness as it limits molecular diffusion towards the growing front [20]. Moreover, contact angle and thermal hysteresis calculations revealed FucoPol indulges in Gibbs-Thomson surface binding, the same adsorption-inhibition mechanism observed for antifreeze proteins [21–24], and often attributed to the presence of hydrophobic groups as well [25–27]. The evidence of Gibbs-Thomson growth modulation corroborated initial findings on the paradoxical anticipation of crystallization [6], but isochoric super-cooling experiments designed to capture a polysaccharide influence on the stochastic phenomenon of nucleation revealed another mystery. Contrary to small molecule cryoprotectants [28], FucoPol generated a drastic narrowing of the nucleation temperature distribution of pure water, an effect magnified by increased FucoPol concentration. In other words, the aqueous FucoPol-water system became increasingly deterministic [29]. Moreover, when nucleation probability was expressed as a function of time (Poisson-distributed), the induction time (*τ*) necessary for the first nuclei to emerge exponentially increased with FucoPol concentration and viscosity but remained unchanged with small cryoprotectants. This is supportive of an anti-nucleation effect, but conflicts with the anticipation of nucleation [29] (higher *T*_*n*_) and crystallization [6] (higher *T*_*c*_) phenomena, which both suggest pro-nucleation effects instead. Despite the contradicting anticipation of nucleation and ice growth, i.e. the promotion of the exact phenomenological effects responsible for cryo-injury, the biological post-thaw benefits of these polysaccharides to cryopreserved cells was undeniable. In fact, the strategy of eliminating supercooling-based extensive ice growth by manually injecting mineral pro-nucleators into microliter systems has shown strong biological benefits [30, 31], but this effect had never been observed with large molecule polysaccharides, which prompted the search for a deeper mechanistic understanding of their function. In this paper, we leverage this compilation of knowledge, which culminated in the mapping of a complex interplay 1 and further discuss this paradoxical duality of simultaneous observations of anti-nucleation (ANTI) and pro-nucleation (PRO) in subzero polysaccharide-water systems. To answer this, we resort to heuristic thermodynamic modelling and dissect the nucleation timeline from different frames of reference, to put forth a unifying hypothesis that may explain the observed phenomenology. Such a type of model would enable predictive screening of new cryoprotective polysaccharide candidates and shed light on the behavioral dynamics of polymeric agents.

## Materials & Methods

### Classical Nucleation Theory (CNT) modelling

The energetic landscapes plotted in Figures 4, 5, 6, 8, and 9 follow the mathematical formalisms of Classical Nucleation Theory (see SI *Chapter 1*). Briefly, the nucleation activation energy barrier Δ*G*_*n*_ is a gain-cost function, expressed as the linear combination of a volumetric free energy gain component (VFE, or Δ*G*_*v*_) arising from agonistic nuclei growth, and an interfacial energy antagonistic cost component (IE, or *γ*_SL_) arising from the repulsive interaction between a frozen fraction surface of angular convexity *θ* and the migration of new water molecules from the bulk into a growing nucleus; and can be expressed as follows:

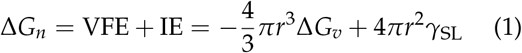

where Δ*G*_*v*_ is the volumetric free energy and *γ*_SL_ is the solid–liquid interfacial energy. The preceding coefficients of both energy parameters correspond to the volume and surface area of a cluster, assuming a perfectly spherical, homogeneously nucleated geometry. By interactive iteration of these formulas, a change in Δ*G*_*v*_ and *γ*_SL_ was performed to obtain extreme–value and optimal energetic landscape scenarios to accommodate the observed phenomenology.

**Figure 1:**
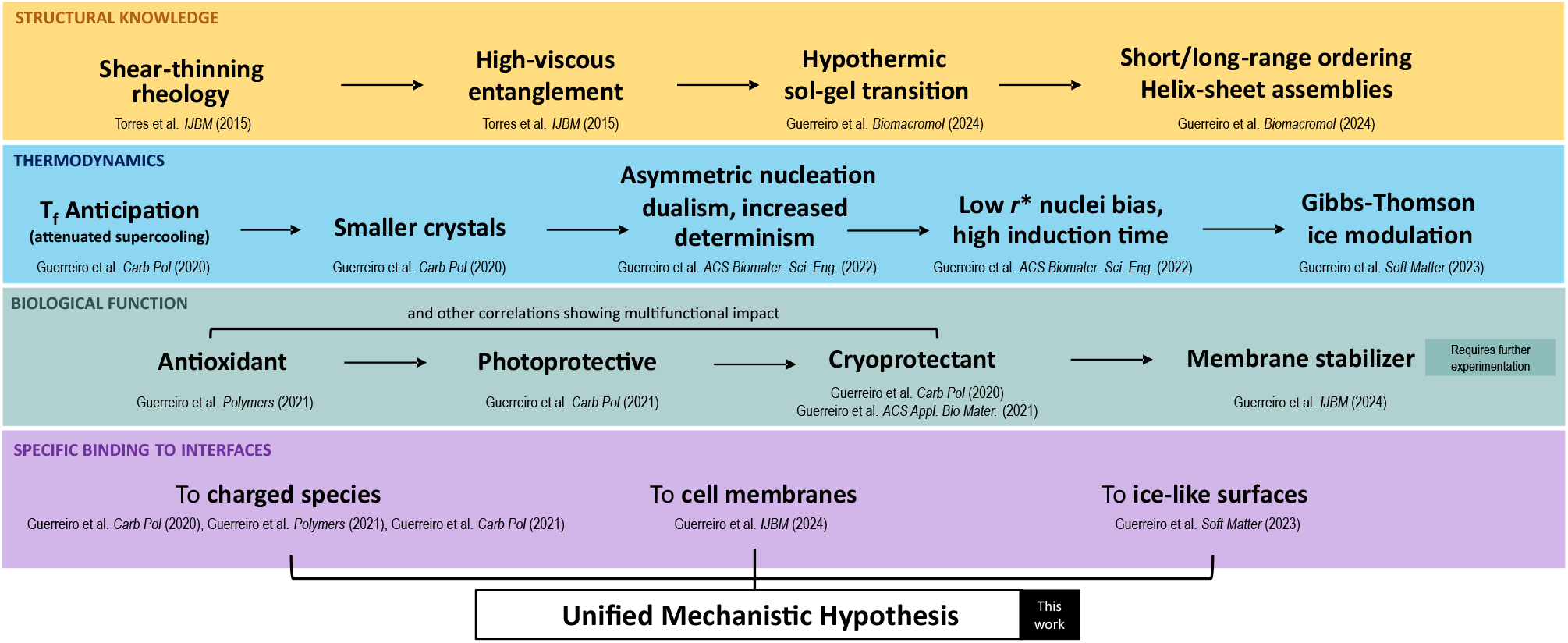
Compilation of structural, function, proprietal and action-by-design characteristics of cryoprotective polysaccharides published previously. The interplay between these traits provides a structure-function relationship mapping for cryoprotective polysaccharides.

**Figure 2:**
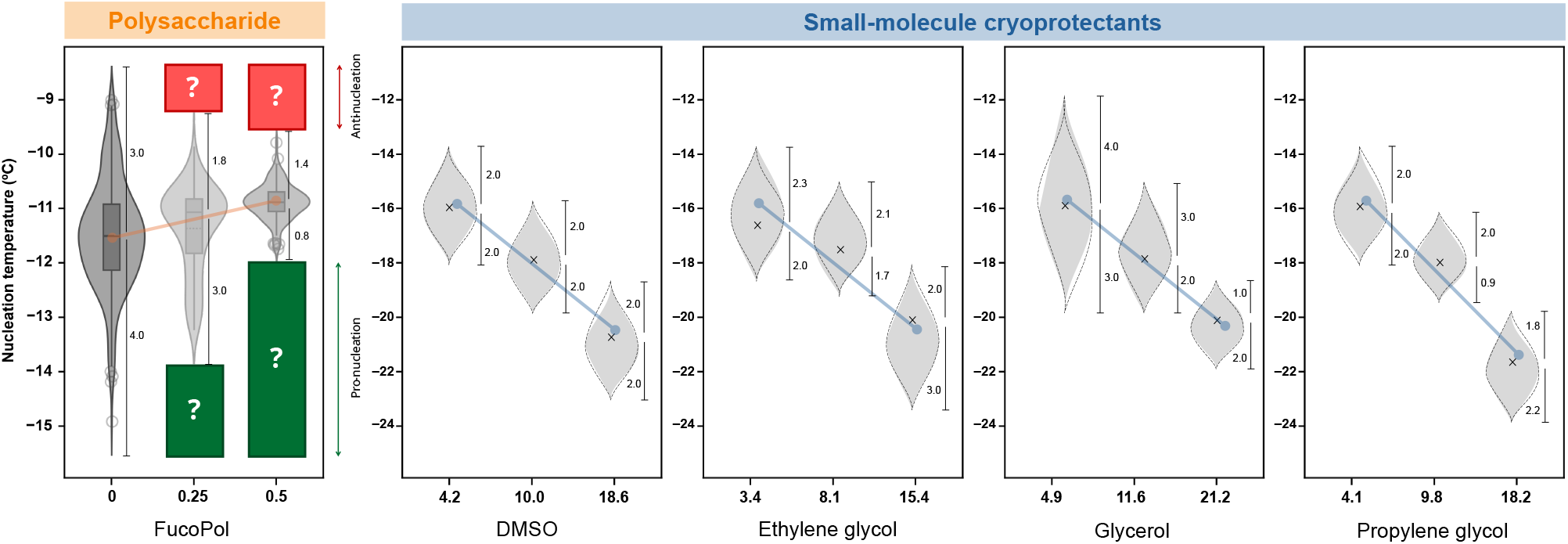
Isochoric supercooling nucleation experiments, contrasting polysaccharide and small-molecular cryoprotectants. (a) Evidence of asymmetric stochastic narrowing observed for FucoPol solutions [29]. Green- and red-shadowed regions represent temperature domains of nucleation extinction and, respectively, suggest pro-nucleation and anti-nucleation phenomena. (b) Comparison with data collected by Consiglio *et al*. [28] on small-molecular cryoprotectants using the same isochoric device, but a different pure water standard. Band width, mean *T*_*n*_, and Δ*T* around mean *T*_*n*_ measured by solid black lines are only approximate representations for the sake of comparison. Band profiles are not true to reality. For the original accurate data collection and analysis, the reader is referred to the original paper [28].

**Figure 3:**
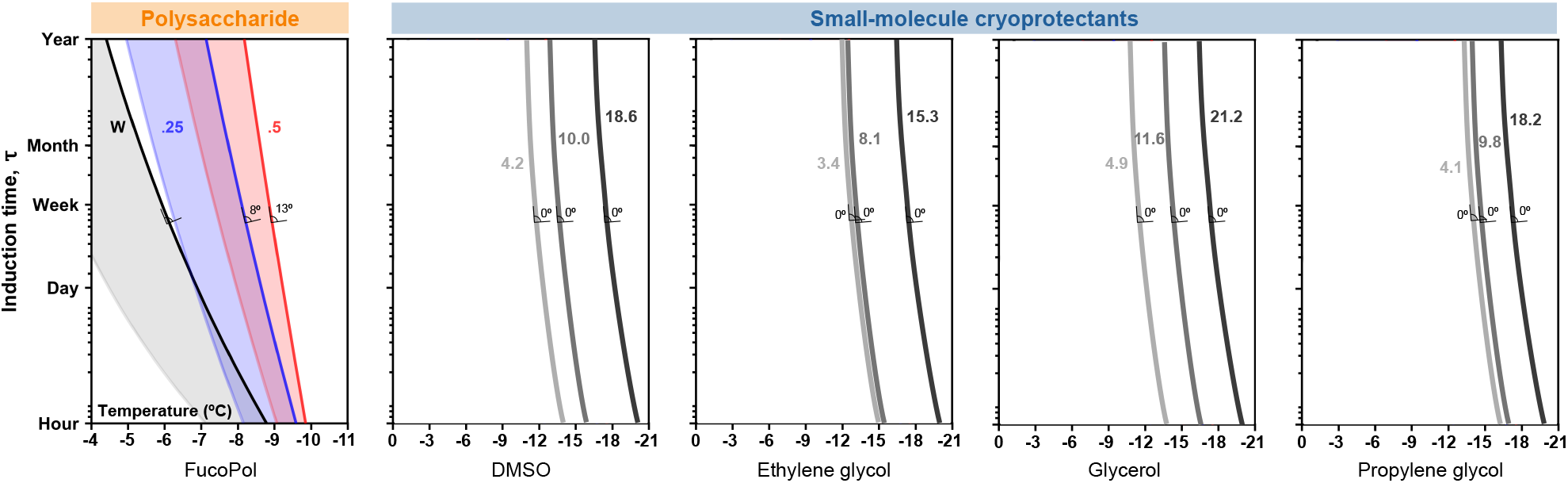
Poisson-modelled induction time bands observed for FucoPol solutions [29] and comparison with data collected by Consiglio et al. [28] on small-molecule cryoprotectants using the same isochoric device, but a different pure water standard. Data labels near curves represents the mol% concentration of each sample. The slope angle for each angle is relative to its pure water sample standard. The x-axis represents system temperature in °C. The *τ*/*T* is a representation of system nucleation susceptibility: at a given *T*, nucleation probability is maximal at time *τ*. FucoPol was the only molecule for which a concentration increase resulted in a change in *τ*. The induction time curves for small-molecule cryoprotectants are only approximate representations for the sake of comparison. The pure water data of Consiglio et al. to which the slope angles refer to, is not shown. For the original accurate data collection and analysis, the reader is prompted to the original paper [28].

**Figure 4:**
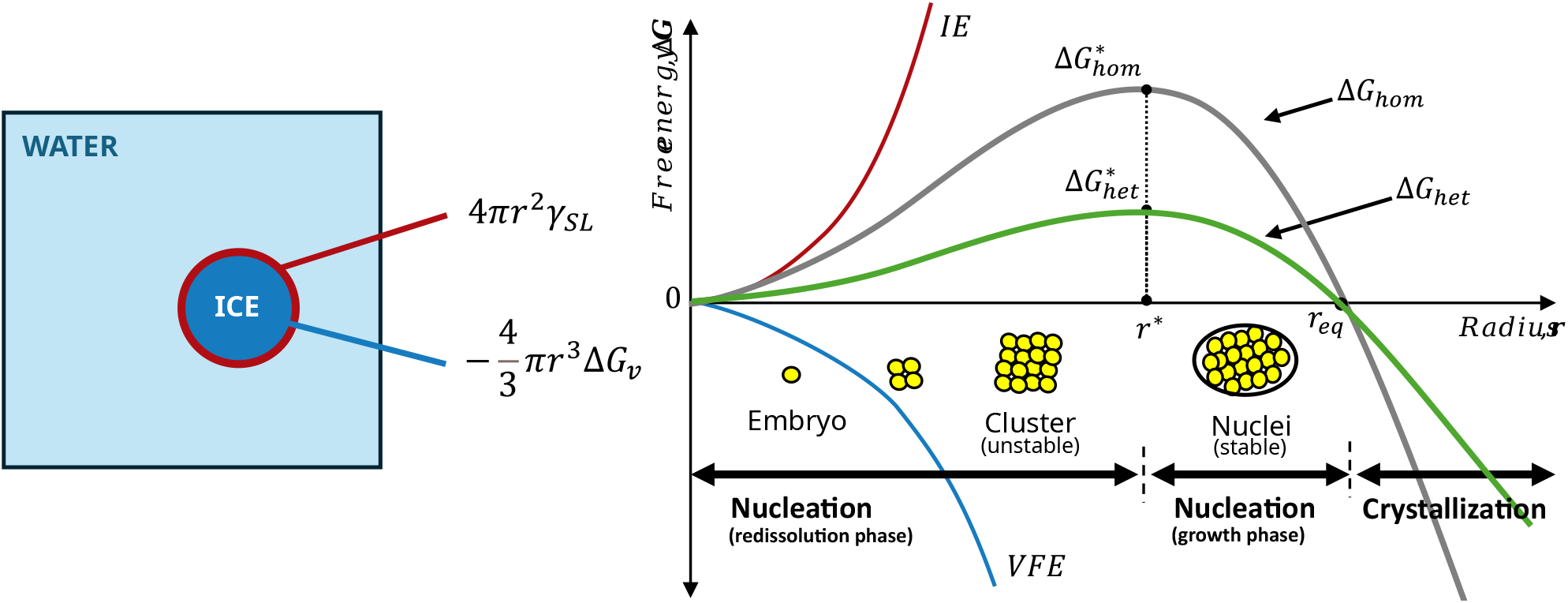
The energetic landscape of nucleation. The onset of crystal growth is a fine equilibrium between interfacial energy (IE, red) and volumetric nuclei morphology factors (VFE, blue). The growth of ice nuclei minimizes the energy required for ice growth to occur because the volumetric free energy scales with system volume and molarity. On the other hand, a bigger nucleus has a greater surface tension due to unfavorable curvature, leading to a required activation energy (green) expressed by Eq. 13 (*SI*). Upon reaching a critical nuclei radius *r*\*, clusters become stable and continuously grow until an equilibrium radius *r*_eq_ is reached, leading to crystallization at Δ*G <* 0.

**Figure 5:**
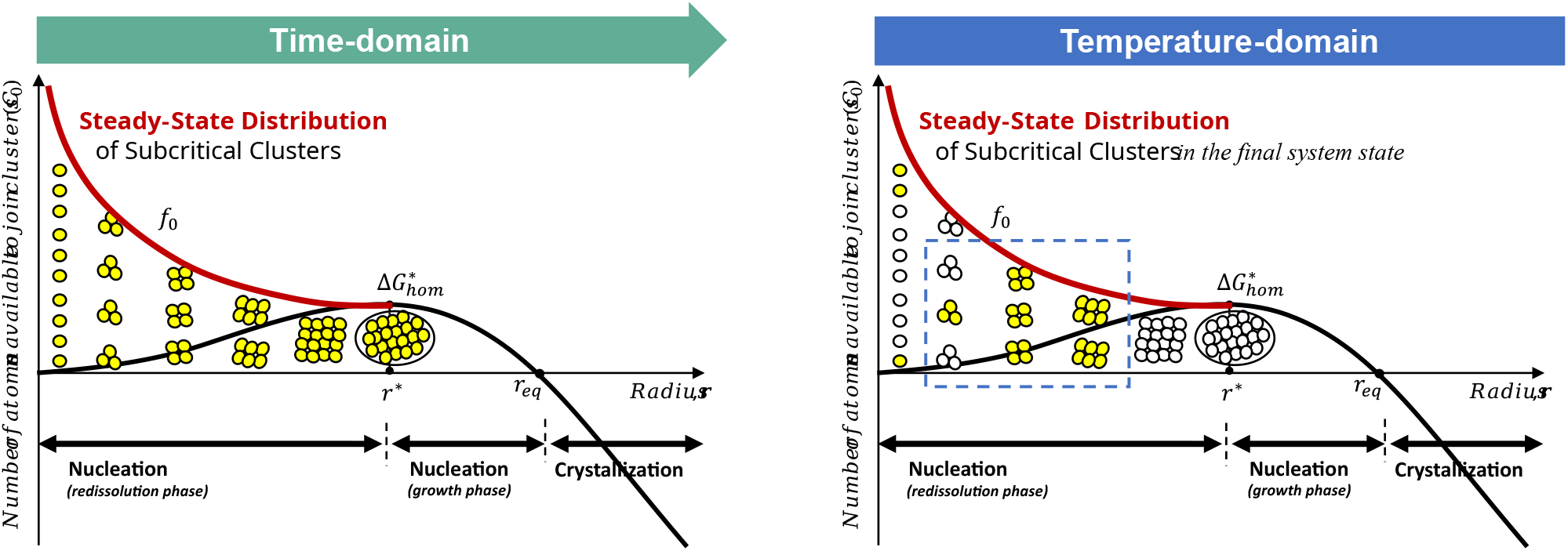
Time and temperature domains used to rationalize nucleation system evolution. The time-domain perspective is predominantly theoretical and rationalizes the gradual growth of nuclei at different stages of their inception, thus populating the whole spectrum of subcritical cluster sizes. The temperature-domain perspective is a practical rationalization of a steady-state growth Boltzmann distribution of cluster sizes that represents a final system state and is sensitive to size constraints imposed by initial system conditions.

**Figure 6:**
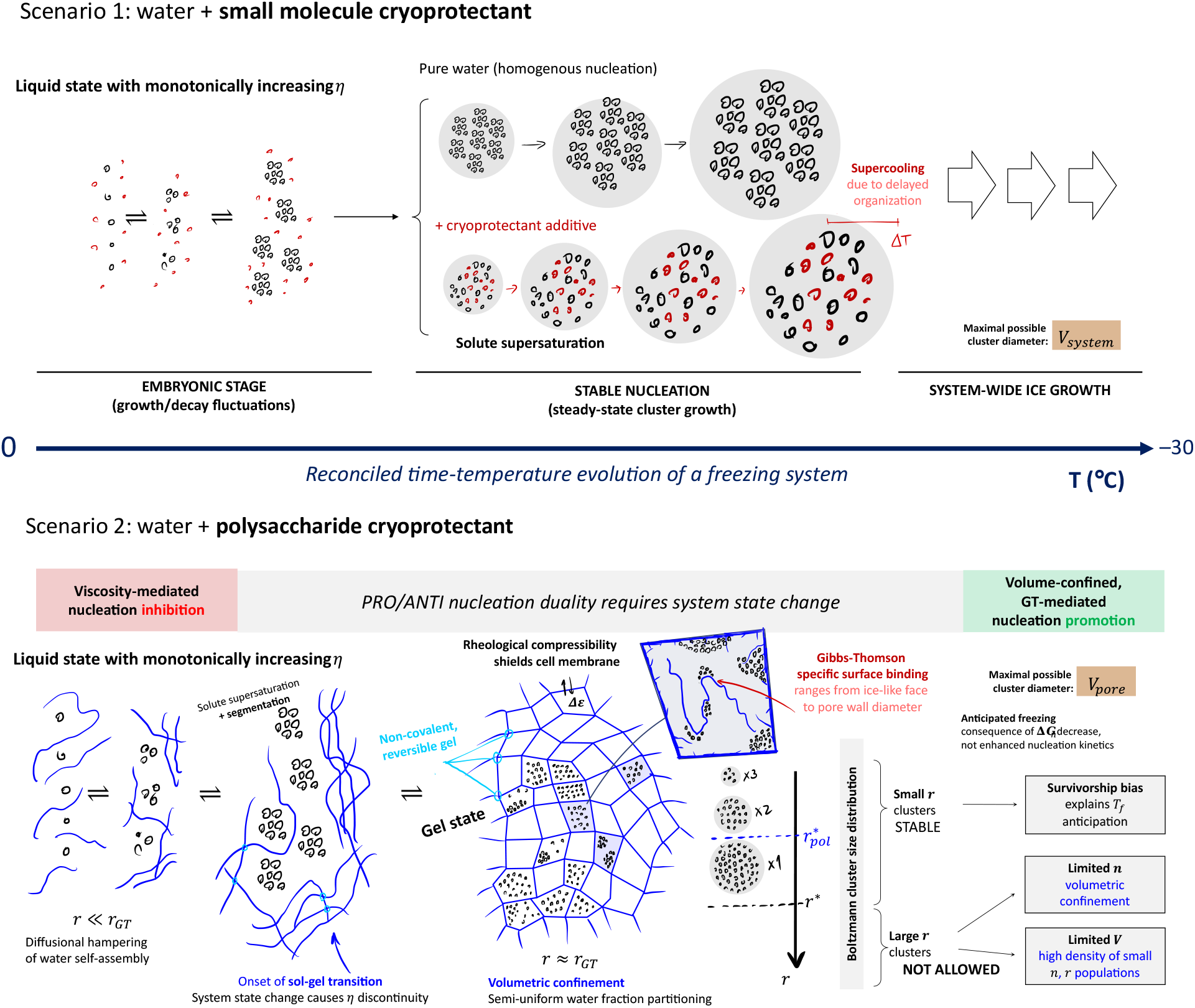
A schematic representation of the rationale behind the unified mechanistic hypothesis for polysaccharide cryoprotection hereby proposed. The contrast between the behavior of monomeric (*scenario 1*) and polymeric (*scenario 2*) cryoprotectants are shown, with particular emphasis on the pore-forming capacity being a critical feature in compartmentalization of unfrozen fractions and subsequent effects on nucleation, also explaining how the nucleation duality may arise.

**Figure 7:**
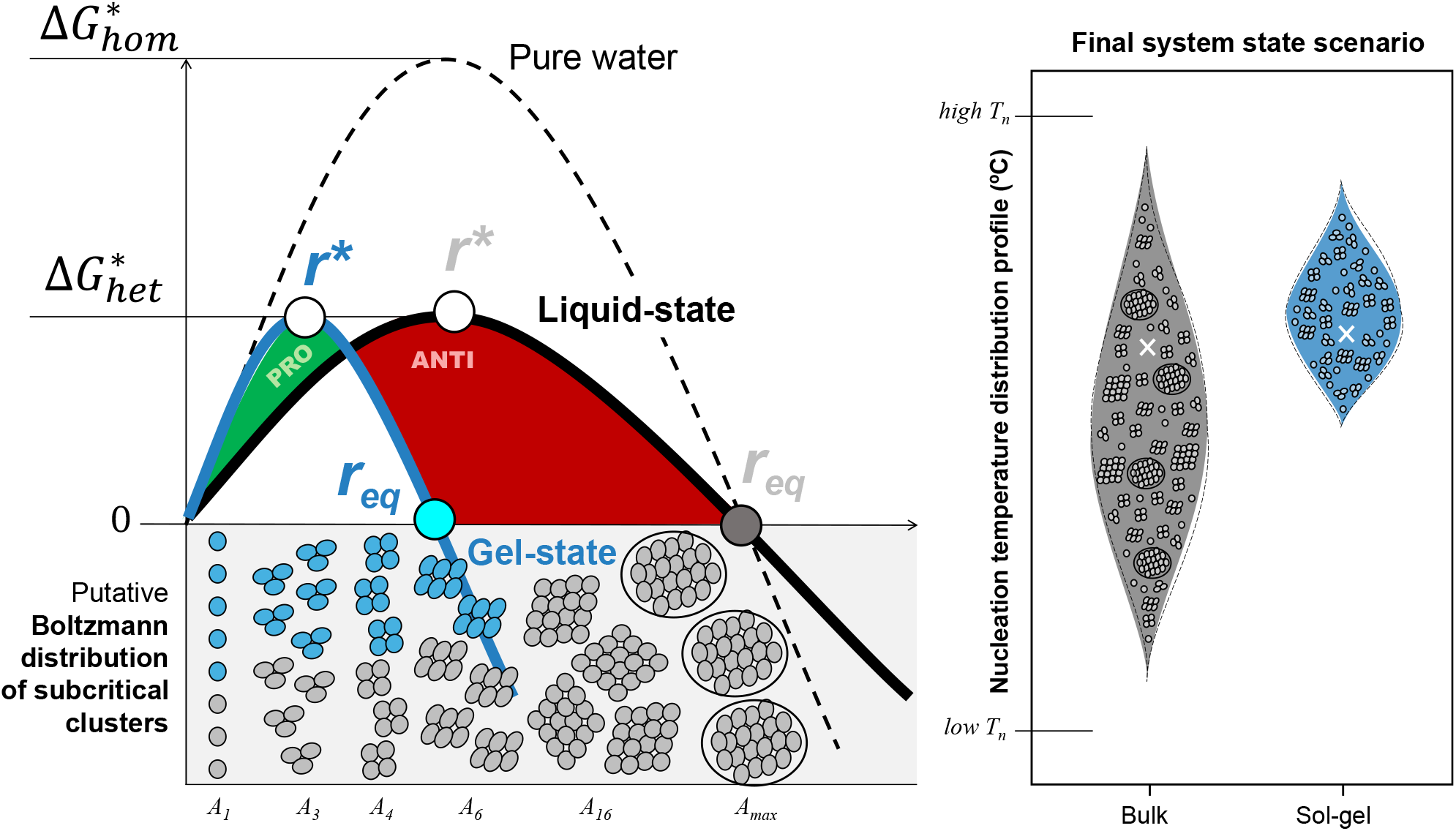
Simulation of the approximate CNT landscape of a system before (grey) and after (blue) a sol-gel transition occurs, highlighting how the generation of compartmentalized gel pockets may influence the VFE and IE energy contributions towards nucleation emergence (left) and the nucleation temperature profiles thereafter (right). After a system state change, the anti-nucleation (red) and pro-nucleation (green) shaded regions emphasize which alterations are observably expected, namely an inhibition of nuclei growth with 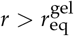 leading to concomitant nucleation promotion at lower 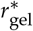 sizes. An illustrative representation of Boltzmann cluster sizes along the *r*-axis and as populations within the system’s stochastic bands is also shown to emphasize the small nuclei survivorship bias observed.

**Figure 8:**
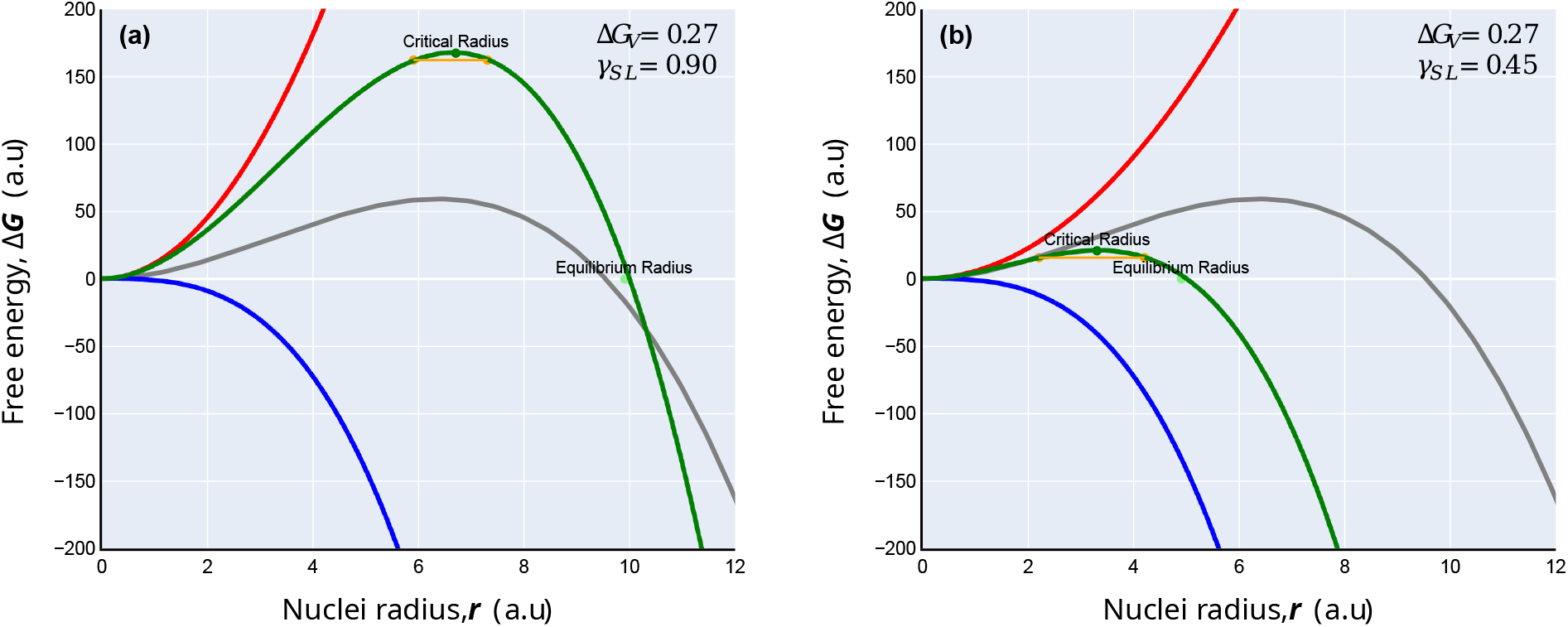
Free energy landscapes of nucleation phenomenology, according to Classical Nucleation Theory. In gray, the theoretical heterogenous nucleation of pure water is shown for reference. A two-fold reduction in *γ* at constant Δ*G*_*v*_ leads to a shift from anti-nucleation (a) to pro-nucleation (b) landscapes.

**Figure 9:**
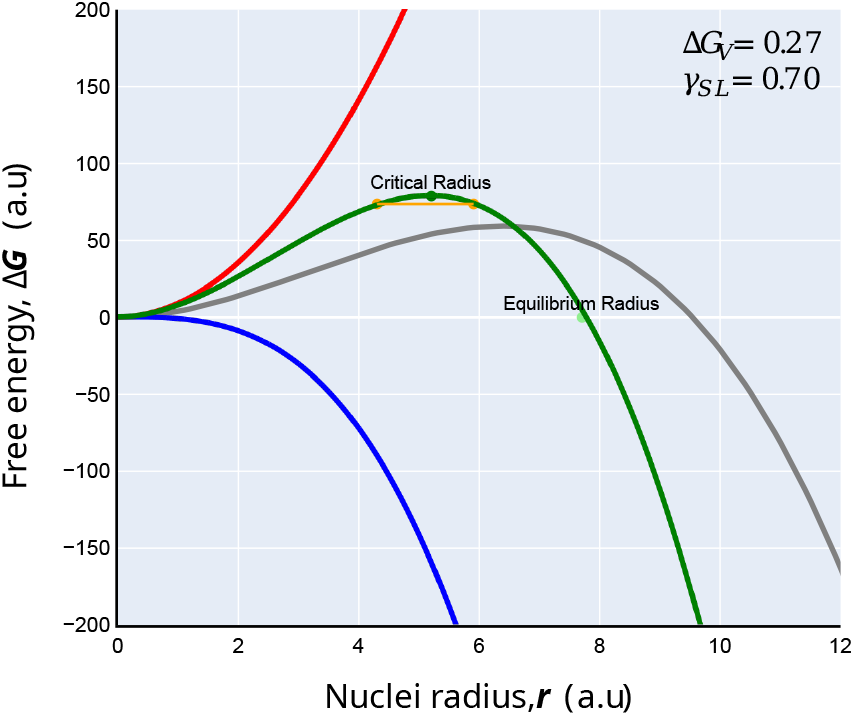
Best-parameter CNT energetic landscape, accounting for Zeldovich instability associated with thermal fluctuations *kT*, iteratively obtained in order to reconcile pro-nucleation and anti-nucleation effects. In gray, the theoretical heterogenous nucleation of pure water is shown for reference.

## Results

Certain particularities in previously collected data prompted a deeper analysis into how the observed phenomenology may be reconciled with known nucleation physics models, but the sparsity of knowledge at the time did not allow it. Now, we recall and perform a critical analysis of several points: the particular asymmetry in the stochastic narrowing of nucleating water in the presence of FucoPol; the root drivers of anti-nucleating and pro-nucleating behavior, their simultaneity and sequentialism; the impact of conformational changes, molecular order-disorder and sol-gel transitions in the thermodynamic potential of nucleating systems; and ultimately the fit between these observations and Classical Nucleation Theory models.

### Asymmetric stochastic narrowing

In isochoric supercooling experiments of FucoPol– water systems, a narrowing of the nucleation temperature band was observed [29]. The stochastic nature of nucleation implies that its phenomenological emergence cannot be predicted at a given *T*, but can be determined to occur in a predictable range Δ*T*. A narrowing of this range implies a system that has become less stochastic, or more deterministic. However, the stochastic narrowing of the *T*_*n*_ distribution was asymmetric and skewed towards high *T*_*n*_ regimes, an observation more evident with increased FucoPol concentration. The mean *T*_*n*_ obtained was strongly correlated with a Gompertz rheological viscosity curve, and deep nucleation temperatures were increasingly extinguished [29]. Figure 2 reiterates the obtained nucleation temperature distribution bands with FucoPol, emphasizing the temperature regions where ice nucleation events became extinct. The most evident difference between FucoPol and the monomeric cryoprotectants surveyed by Consiglio *et al*. [28] was that small molecules result in drastic changes of average *T*_*n*_ but do not generate a consistent asymmetric stochastic narrowing, contrary to the polysaccharide. The extinction of high *T*_*n*_ regimes (Figure 2, red) is indicative of an anti-nucleation effect that is expected from any cryoprotectant which leads to ice growth inhibition by supercooling the system. This reflects the observed behavior for most cryoprotectants, as observed for DMSO, ethylene glycol, glycerol, and propylene glycol, and is most likely associated with kinetic inhibition, as the extinction is magnified with increasing concentration (and therefore, viscosity). A pro-nucleation effect indicating an anticipation of the nucleation event would in turn result in an observable population of the upper distribution tail (above average *T*_*n*_), not its extinction. But this was not observed; rather, the extinction of low nucleation regimes (below average *T*_*n*_) on the other end of the distribution was visible. Although this also remains an indicator of pro-nucleation, it implies a density increase of events near *T*_*n*_ to sustain the principle of mass conservation. Anticipated nucleation in the temperature scale would provide the initial conditions for a subsequent anticipation of crystallization and reduction in crystal size dimension, as observed before [6], without requiring accelerated growth kinetics in an identical time frame.

Therefore, the behavior of FucoPol suggests a unique thermodynamic ANTI/PRO nucleation duality, where there exists a paradoxical, simultaneous expression of anti-nucleation and pro-nucleation. Thermodynamically, the stochastic narrowing being asymmetric suggests that a nuclei survivorship bias exists predominantly towards promoting the emergence of small-sized nuclei early on, with complete inhibition of nucleation events at lower temperatures. However, the opposite was inferred in induction time *τ* studies as a probability of nucleation emergence (Figure 3). Consiglio et al. [28] noticed that small molecule cryoprotectants consistently supercooled the system with no change in nucleation susceptibility (constant *τ*), but FucoPol actively demonstrated a stabilization of the supercooled system. An exponential increase in *τ* at high *T*_*n*_ is suggestive of anti-nucleation due to a drastic increase in the nucleation energy barrier. However, an increasingly deterministic system also resulted in a steeper *τ*/*T*, which suggests that discrete temperature changes at lower *T*_*n*_ results in greater nucleation susceptibility, hence pro-nucleation. The isochoric supercooling experiment allowed to dissect nucleation dynamics from temperature (Figure 2) and time (Figure 3) domains. From nucleation temperature distributions pro-nucleation appears dominant, but an increased induction time until the first nuclei emerge is suggestive of anti-nucleation. In both analyses, the duality still holds, but on account of which phenomenon arises first, they led to antagonistic conclusions for the influence of FucoPol.

Hence, from this apparent dual behavior, two questions arise:

1. What thermodynamic rationale allows to reconcile the fact that both pro-nucleation and antinucleation may occur in the system?
2. Is there a causal link between pro-nucleation and anti-nucleation, as in one phenomenon resulting in the expression of the other? And if so, by which order is that temporal causality true?

To answer these questions, we deconstructed the observed phenomenology from basic thermodynamic and kinetic assumptions (SI *Chapter 1*) and rationalized the nucleation event from temperature-domain and time-domain perspectives.

### Causality and sequentialism in nucleation phenomenology

The nuclei survivorship bias hypothesis supporting pro-nucleation was reasoned to be valid from first-principle properties of thermodynamic systems with limited mass amount. The Helmholtz thermodynamic potential in isochoric systems states that, in a constant-volume recipient:

i. There exists a finite amount of liquid water which can nucleate.
ii. Extensive nucleation early on reduces late-stage nucleation probability due to a reduction in chemical potential.
iii. Homogenous nucleation from novel nuclei is energetically less favorable than heterogenous nucleation from pre-existing clusters, thus less likely.

First, these properties assume an implicit causal relationship, where early events and initial conditions determine the final state of the system. Second, the emergence of new nuclei always requires overcoming an interfacial energy cost (Figure 4) far greater than that required for stable clusters with radius already larger than their critical radius (*r*\*) to grow. This cost disparity is magnified (i) at lower temperatures and (ii) in increased viscosity environments, both due to reduced molecular mobility. As thermodynamic systems evolve by minimizing their variation in internal energy, the generation of new nuclei at low *T*_*n*_ has very low likelihood, thus corroborating the low *T*_*n*_ extinction observed (Figure 2, green). However, a causal relationship also implies that this extinction is induced by pro-nucleation towards high *T*_*n*_ because both effects are related by resource extinction: the lack of freely available water molecules with high diffusivity at lower *T*_*n*_ reduces the chemical potential for nucleation; but an extensive nucleation at high *T*_*n*_ would also result in a reduced unfrozen water fraction at low *T*_*n*_, leading to the same system state. The issue with heuristic rationalizations of causality is that pinpointing a linear sequence of events varies according to the frame of reference. If the pro-nucleation effect of FucoPol is dominant due to a greater bias in asymmetric narrowing towards high *T*_*n*_ events, why does an exponential increase in *τ* indicative of stabilized ice-free supercooled regimes largely supports anti-nucleation? Second, if one considers the temporal evolution of a system initially at 0*°*C being continuously cooled, then would not the onset extinction of high *T*_*n*_ values between *−*7*°*C and *−*9*°*C be indicative of an anti-nucleation effect occurring first instead? While these paradoxical observations do not disprove the existence of a duality, they highlight that an ANTI/PRO behavioral shift must exist in a linear timescale.

### Referential frames for system evolution: time, temperature

To rationalize this ANTI/PRO nucleation duality and its causal link, the progressive evolution of a nucleating system was interpreted in terms of temperature variations at constant time, *t* (temperature-domain, *∂T*); and in terms of elapsed time at constant temperature, *T* (time-domain, *∂t*). These are heuristic interpretations (Figure 5), as it is evident that during cooling, *∂T* and *∂t* are co-variant, but they help in rationalizing why the observed *T*_*n*_ distributions in Figure 2 do not reflect a logical temporal evolution of the system. From a time-domain perspective, a single freeze-thaw cycle may be interpreted as a linear sequence of events. At constant *T*, the thermodynamic system can still evolve because nucleation is a stochastic event with an associated time-bound Poisson probability (see *SI* and Figure 3). A supercooled system may be held at constant *T* for an undefined period of time, for which there is an associated probability of nucleation occurrence [29, 32]. However, the ability to be informed for how long a supercooled system may remain ice-free at a given *T* does not mean the system is nuclei-free, as they may grow over time until *r*\* is reached. From a temperature-domain perspective, the characteristic nucleation distributions obtained (Figure 2) are a final representation of system state. Rather than informing about system evolution, they are ensemble representations of the behavior of a thermodynamic system, a statistical approximation that originated from numerous freeze-thaw cycles where an observed stochastic emergence of nucleation was recorded. So, a final system state is a Boltzmann population distribution profile of nuclei sizes at a given temperature, assuming a steady-state growth rate of the subcritical clusters, such as the one represented in Figure 5. Assuming an arbitrary water cluster *A*, containing *n* water molecules, of size *A*_*n*_, it is expected that at the onset of cooling, only small cluster sizes e.g. *C*(*A*_1_), *C*(*A*_3_) may be fully populated by recorded nucleation events, where *C* is the amount of clusters of size *A*_*n*_ according to the steady-state distribution defined by Eq. 25 (*SI*). As temperature decreases, the system approximates to a normally distributed cluster population, from *C*(*A*_1_) up to a size where the critical *r*\* condition is satisfied. Above *r*\*, stable nuclei will not re-dissolve, continuously growing until reaching *r*_eq_ and leading to spontaneous ice growth. While temperature-domain interpretations lead to inferring a simultaneous emergence of anti-nucleation and pro-nucleation effects in the final system state, this emergent duality is nonsensical from a time-domain perspective. For instance, the increased viscosity of a water-FucoPol system during the early stage of cooling results in reduced water molecule mobility and consequent diffusion to growing clusters. Therefore, the first observable phenomenon would be a kinetic delay in nucleation relative to pure water systems, resulting in anti-nucleation. This is supported by the observed high *T*_*n*_ extinction (Figure 2, red) and higher *τ* (Figure 3). Viscosity monotonically increases during cooling according to a rheological Gompertz growth model [29], so the established anti-nucleation effect should be preserved and further magnified. However, Figure 2 shows a predominant extinction of low *T*_*n*_ events, which by analogy with Boltzmann distribution profiles (Figure 5), indicates an extinction of large clusters e.g. *C*(*A*_40_) in favor of small clusters e.g. *C*(*A*_1_*−*_7_), which is suggestive of pro-nucleation. In the time-domain analogy, this simultaneity is non-sensical as one effect must predominate. In light of this paradox, the simultaneous emergence of opposite nucleation phenomena does not seem possible in an unchanged chemical system, and a rationalization of the cooling phase in the same nucleating system using different heuristic models elicits a causal loop.

Therefore, we have conceived that this duality is only feasible if two events undeniably occur:

1. There must exist a discrete turnover point between system states, over time, which allows for pro-nucleation to dominate over anti-nucleation.
2. There must exist a physicochemical change in system character that enables the behavioral shift.

### The unified mechanistic hypothesis

All previous rationalizations argued that a causal link must exist, but none truly explained (i) the propensity towards FucoPol being a pro-nucleator (although collected data consistently suggests so [6, 29]), (ii) what system change drives an ANTI/PRO behavioral shift, or (iii) the reasoning behind an increased determinism of a phenomenon which, by nature, is intrinsically stochastic. In a past structure-function relationship study that screened biologically cryoprotective and non-cryoprotective polysaccharides [4], those who elicited attenuated supercooling – the anticipation of crystallization relative to supercooled pure water – expressed the highest degree of biological cryoprotection. The withstanding explanation at the time was that precocious ice growth within the same time frame led to the formation of stable crystals so small that effectively they were innocuous to biologicals [6]. The initial assumption was an acceleration of growth kinetics, but the Gibbs-Thomson growth modulation observed in directional freezing experiments [20], which resembled the mechanism of an antifreeze protein, and the increased *τ* characteristic of supercooled system stabilization [29] both hinted at an inhibitory effect based on system energetics, hence thermodynamic in nature. In that same study [4], attenuated supercooling was accompanied by two critical observations: (i) that polysaccharides with biological cryoprotection traits also possessed the ability to structurally assemble into conformations with a higher degree of order than random coils; and that (ii) this would potentiate a reversible sol-gel transition in hypothermic regimes. The discovery of a mechanism resembling reversible cold gelation in cryoprotective polysaccharides could support the necessary evidence for a change in system character that would originate an ANTI/PRO behavioral shift. Therefore, we put forth in this unified hypothesis that the cryoprotective mechanism of polysaccharides arises from a combination of three features (Table 1), without any of which the theory fails to reconcile the nucleation duality with the accumulated cryoprotective evidence so far. Polysaccharides that show cryoprotective traits by anticipating nucleation and ice growth may show a combination of (i) kinetic hampering of molecular diffusion (concentration effect), (ii) Gibbs-Thomson specific ice binding (templating effect) that leads to growth modulation and size reduction, and (iii) the formation of a gel architecture of defined porosity (mesh size effect) that explains how a system initially anti-nucleating may undergo an ANTI/PRO behavioral shift amid cooling to become predominantly pro-nucleating, leading to an increased stochastic determinism. We further advance, in retrospective, that attenuated supercooling is not the originating phenomenon of early ice nucleation/growth, but rather a consequence of the polysaccharide matrix transitioning into a gel state, whose new features may propitiate such an anticipation in phenomenology. The generation of a porous architecture through self-assembly amid system cooling not only provides a system state change to explain the behavioral shift but also supports a system disposition towards the observed nuclei survivorship bias, the increased stochastic determinism, and the pro-nucleation character. All these hint at a perceived kinetic acceleration, but this causation is *illusive*. Instead, we hypothesize that causality arises from an increase in thermodynamic probability, which in turn is expressed in the form of nucleation anticipation, rather than a direct enhancement of nucleation kinetics. We shall now proceed to examine the claims of our hypothesis in greater depth.

**Table 1:**
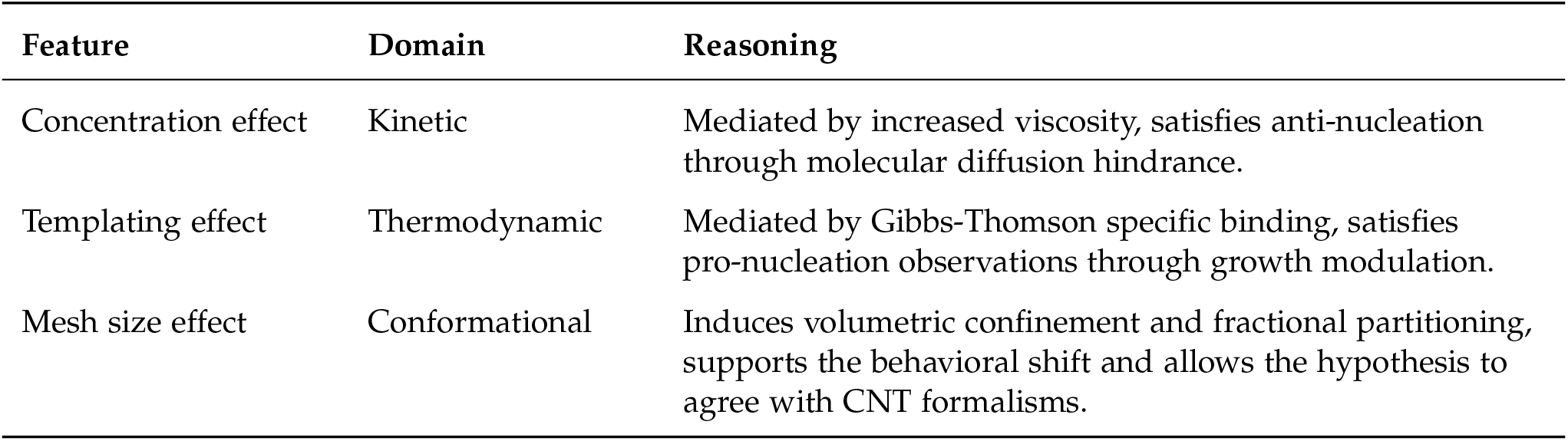
Rationale of the unified mechanistic hypothesis in cryoprotective polysaccharides.

| Feature | Domain | Reasoning |
| --- | --- | --- |
| Concentration effect | Kinetic | Mediated by increased viscosity, satisfies anti-nucleation through molecular diffusion hindrance. |
| Templating effect | Thermodynamic | Mediated by Gibbs-Thomson specific binding, satisfies pro-nucleation observations through growth modulation. |
| Mesh size effect | Conformational | Induces volumetric confinement and fractional partitioning, supports the behavioral shift and allows the hypothesis to agree with CNT formalisms. |

### Claim #1: Volumetric confinement and fractional partitioning

Figure 6 shows a schematic representation of the unified mechanism hereby proposed, from a temperature evolution perspective. In the initial stage of cooling, small molecule cryoprotectant systems are virtually identical to polymeric systems in what concerns water molecule aggregation into smaller nuclei, or embryos. Both systems undergo an increase in viscosity, but polysaccharides accentuate this increase, leading to earlier and greater loss of molecular mobility. A continuous decrease in temperature leads to decreased collision frequency and attractive forces will dominate over repulsive forces, resulting in clustering. As the unfrozen fraction further reduces due to water migration towards cluster growth (increasing *r*), solute supersaturation further leads to nucleation inhibition by kinetic hindrance. In the case of small molecule cryoprotectant systems such as glycerol and DMSO, the disruption of water molecule directionality delays the periodic organization of embryos into larger stable nuclei near *r*\*, thereby resulting in system supercooling and ice growth inhibition (Figure 6, scenario 1). However, polymeric systems contain an additional source of attractive forces, namely chain-chain interactions, which largely contribute to polymer entanglement. If this conformational arrangement is disordered, polymer chains largely increase solution viscosity and further indulge in kinetic inhibition: promoting anti-nucleation. This is corroborated by high *T*_*n*_ extinction (Figure 2) and increased *τ* (Figure 3). As previously mentioned, cryoprotective polysaccharides which demonstrated an ordered self-assembly also showed the ability to form a gel state, and subsequent attenuated supercooling [4]. The plausibility of a gelling polysaccharide being capable of establishing a porous matrix which confines the unfrozen water fraction into small, compartmentalized pockets (Figure 6, scenario 2) raises major implications in nucleation outcome. Nucleation is largely reliant and very sensitive to the initial conditions of a cooling system and draws its evolution from chaos theory models due to its stochastic nature. If a sol-gel transition occurs and a porous matrix is formed, the unfrozen water fraction which was available to nucleate in a bulk environment is now enclosed in small-volume gel pockets, and a change in system state has occurred. Therefore, the initial conditions for nucleation largely shift mid-cooling, and a new set of circumstances may lead a system which was largely anti-nucleating to behave as pro-nucleating, or *vice versa*. In this case, one can argue that each pore is now its own nucleating system with a unique set of initial conditions for cooling that differ from the bulk. Moreover, if during formation, the gel pores are equalized enough in terms of dimensions and unfrozen fraction volume, then the breadth of possible initial conditions is narrow enough to generate stochastic narrowing. In other words, greater pore uniformity leads to very similar initial conditions, which in turn increases nucleation determinism. From this perspective, the extensive evidence of small cryoprotectants not influencing the stochastic range of nucleation temperatures as observed by contrast in Figure 2 and Figure 3 arises from their inability of undergoing a system state change, e.g. by means of pore formation, like cold-gelation polysaccharides do, regardless of their ability of extensively increasing viscosity. Thus, the nucleation profiles of small-cryoprotectant systems still resemble their pure water-like nature, because all of them still permit bulk nucleation. Nucleation depends on extensive variables such as global system volume *V* and number of water molecules *n* available to join the growing clusters. So, the confinement of water into (*V*/*n*) pockets satisfies the observations of reduced nucleation stochasticity [29] and the consequent reduced crystal size dimensions with a narrowed size distribution [6]. We expect this effect to be magnified by increased pocket size uniformity. This conformational mesh size effect stands as the key feature in this hypothesis for two reasons. First, the only reasoning currently supporting increased stochastic determinism is the uniformization of the initial conditions for nucleation by fractional partitioning of the system bulk into compartmentalized pores. Second, the survivorship bias towards smaller *r* nuclei is only congruent with an existing volumetric confinement that limits the maximal *n* able to migrate to clusters.

### Claim #2: Attenuated supercooling as a consequence of confinement

From a perspective of system energetics, consider a single, confined-volume, gel pocket saturated with water molecules. For simplicity, this unit is considered adiabatic (no heat or mass transfer) relative to neighboring pockets in the grid. As the system cools down to sub-zero temperatures, nucleation probability increases accordingly, materializing in a defined stochastic interval Δ*T*_*n*_. CNT predicts that an anticipation of nucleation relative to pure water dynamics in an identical time interval must be accompanied by a decrease in *r*\*, *r*_eq_, or 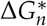 (Figure 7). In a gel pocket, the volumetric free energy contribution is much smaller, so for the average *T*_*n*_ to remain unchanged relative to its distribution span (observed only for the polysaccharide in Figure 2), the interfacial energy must decrease proportionally. So, for a scenario of anticipation nucleation (which precedes attenuated supercooling) to emerge, one of two events must occur:

1. A reduction in ice-water interfacial tension, driven by an ice modulatory effect.
2. An increase in the nucleation rate *J* at *r < r*\* subcritical regimes.

In other words, either the attenuated supercooling effect has a (a) thermodynamic or (b) kinetic origin. Case (a) is plausible from previous data on directional freezing, because FucoPol demonstrated an ice modulatory effect similar to an antifreeze protein, thus it reduces the interfacial energy at the polysaccharide-ice facet [20]. In case (b), *J* may decrease due to viscosity-mediated kinetic inhibition. However, increased viscosity may not necessarily imply a direct influence in *J*, because a kinetic process may be delayed in time but retain its normal rate once thermodynamically permissible. Also, an expressed final effect of attenuated supercooling would imply that, if any change in *J* occurred, it was one of rate increase, not otherwise. Therefore, the thermodynamic origin (a) for the emergence of attenuated supercooling not only is more likely, but it intrinsically establishes that a gel transition should occur, as it requires a confined-volume system state to (i) constrain the maximal achievable *r*\* such that a nucleation anticipation is coupled to a subcritical nuclei population rebalance towards smaller sizes (Figure 7) and (ii) shift from inhibition (anti-nucleation) to enhancement (pro-nucleation). This putative shift in system energetics towards smaller *r*\* and *r*_eq_ would viably represent a new nucleating system composed of nucleation and ice growth anticipation by architectural constraint.

### Claim #3: Gibbs-Thomson templating effect

Volumetric confinement can explain the stochastic narrowing through the assumption of grid size uniformity originating increased determinism; but an asymmetric skewness of the distribution profile towards high *T*_*n*_ regimes implies an active modulation of the interfacial energy between the pocket wall (polysaccharide) and the unfrozen fraction (water) towards favoring pro-nucleation. Therefore, the formation of a FucoPol gel matrix as represented in Figure 6 (scenario 2) suggests the existence of a Gibbs-Thomson templating effect. According to the Gibbs-Thomson (GT) specific ice binding hypothesis [33], a pro-nucleation effect is observed when *r* = *r*_GT_, i.e. the surface radius of a growing ice nuclei (*r*) equals that of a heterogeneous nucleating surface (*r*_GT_) containing an ice-like facet capable of geometrically mimicking the other half of the spherical cluster and significantly reducing the 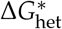 energy barrier. In the absence of a polysaccharide gel matrix, the heterogeneous nucleator is effectively an uncoiled chain where *r≪ r*_GT_, so without specific surface binding, the dominant effect is viscosity-limited anti-nucleation. The assembly of polysaccharide chains into a gel increases the probability for a templating effect to occur, because the effective heterogeneous nucleating surface is now the confining pocket wall, which depending on mesh size, may decrease *r*_GT_ to an extent that the *r* = *r*_GT_ condition is satisfied, and a pro-nucleation effect emerges. This templating effect is as much pronounced as the increase in the ratio between polysaccharide-water interactions and water-water interactions. In gel architectures, the former is expected to drastically increase, in comparison to bulk nucleating systems, where the latter dominates. The predominant stabilization of ice nuclei at smaller *r*\* than pure water, along a limited amount *n* of water molecules in each of the (*V*/*n*) pores, justifies the observed extinction of larger *r*\* population clusters at lower temperatures. Recall the postulation that attenuated supercooling was not a consequence of enhanced kinetics but rather permissive thermodynamics. From a system energetics perspective, this means the probability of nucleation events at higher *T*_*n*_ is increased due to lower 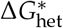, not higher *J*_het_. However, according to CNT [34], pore confinement alone does not automatically bring about a pro-nucleation effect because Δ*G*_*v*_ ∝ *− n*, i.e. the volumetric free energy component that contributes to a decrease in 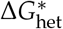 and facilitates nucleation is proportional to the amount of water molecules available to migrate to growing nuclei. Under volumetric confinement, *n*_pore_*≪ n*_bulk_ implies a decrease in nucleation probability, preserving an anti-nucleation effect. Therefore, the existence of a Gibbs-Thomson binding effect is crucial in explaining a minimization in nucleation energetics even under a sol-gel transition. Ultimately, this combination in system conditions appears to be a unique property of gel-forming, ice-binding polysaccharides. Small molecules may be able to form gel-like viscous environments, such as the vitrification of glycerol, but the lack of pore structuring limits the system to only experiencing kinetic inhibition. Gel-forming polysaccharides without an ice-binding response would probably also not elicit pro-nucleation. In summary, these templating and mesh size effects appear to be mutually reinforcing: without pore formation, a strong ice-binding effect would not emerge because *r≪ r*_GT_ would be a constant property of the system; but in spite of pore formation, the cryoprotectant molecule must have the chemical configuration that enables ice modulation in order to exert pro-nucleation, and the dualistic nature observed.

### Re-mapping duality CNT energetic landscapes results in discontinuity

Thus far, we have heuristically demonstrated that a conformational change in system character justifies a change in system energetics. To try reconciling the dual ANTI/PRO phenomenology into a single CNT formalism, we have explored several nucleation energetic landscapes by plotting the CNT equations. These approximate iterations are useful to quickly assess the validity of a hypothetical system without needing to mathematically formalize the thermodynamic constraints that yield a given energetic landscape. Figure 8 therefore represents an extreme-value heuristic modelling of the interfacial energy *γ*, the volumetric free energy Δ*G*_*v*_ and their concerted influence in the resulting energy barrier for nucleation 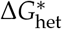. Figure 8a represents anti-nucleation, due to a higher energy barrier 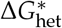 relative to homogenous nucleation, or pure water. Figure 8b represents a scenario where pro-nucleation dominates, achieved here by – all other parameters being kept constant – decreasing by 2-fold the interfacial tension *γ* of the nucleating system. These are extreme-value scenarios where only a single behavior system emerged. The difficulty in justifying a dual nucleation behavior is that, by invoking CNT formalisms, a single energetic landscape could not validly accommodate both effects despite testing a continuum of variable combinations. For a constant-volume system characterized by constant Δ*G*_*v*_, high *γ* implies an increased difficulty in nucleation occurrence, due to an increased nucleation energy barrier (Figure 8a). Under Gibbs-Thomson templating effects, a significant reduction in *γ* due to surface binding cooperativity between the surface of a growing nuclei and the ice-like facet of the polysaccharide structure is expected whenever *r* = *r*_GT_ (Figure *SI*.*2*). A nucleating system appears highly sensitive to shifts in energy gains or costs, such that a 2-fold reduction in *γ* leads to a 7-fold reduction in 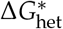 (Figure 8b). Nuclei size variables *r*\* and *r*_eq_ would also decrease. The yellow threshold in Figure 8 centered around *r*\* and of height *kT* represents the Zeldovich factor (*Z*), a thermal metric of nucleation susceptibility (Eq. 19, *SI*). A flatter band centered at 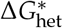 implies a greater susceptibility for nuclei *r ≈r*\* to become stable and continuously grow, because the required thermal fluctuations *kT* near 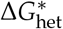 are smaller. All these changes would be indicative of a pro-nucleation effect, relative to pure water conditions. However, the emergence of volumetric confinement contradicts this tendency. A reduction in *n* leads to a decrease in *γ* by a factor of *n*^2/3^, which does not outweigh the *−n* change in Δ*G*_*v*_ (Figure *SI*.*1*). In other words, a drastic decrease in volumetric free energy from pore formation must be outweighed by a balancing act that increases nucleation probability. So, for a predominantly pro-nucleating system to ultimately survive, this heuristic demonstration reinforces that a Gibbs-Thomson ice-binding effect is necessary to drive a reduction in *γ*, even in volumetrically confined systems. By fine-tuning *γ* and Δ*G*_*v*_ one arrives at the most-probable energetic landscape that could accommodate an ANTI/PRO nucleation duality observed for the FucoPol polysaccharide-water system (Figure 9). The energetic landscape depicted conforms to an anti-nucleation effect due to a greater nucleation barrier 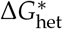 relative to pure water. However, it also implies that its corresponding *r*\* is lower than homogeneous nuclei, an indication of pro-nucleation. The evolution of such a system is physically inconsistent with mathematical CNT formalisms, because an increase in *r*\* along the 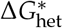 curve (Figure 9) encounters an anti-nucleating regime but the final system state generates *r*\* distributions characteristic of pro-nucleating regimes. Also, the convexity difference between *Z*_het_ and *Z*_hom_ seems insufficient to justify an increased nucleation susceptibility. This inability to simultaneously observe a dual nucleation behavior in a single mathematical CNT landscape reinforces the need to have a change in system state (i.e. a sol-gel transition) to accommodate the physical emergence of both behaviors. Therefore, the causal evolution of nucleation phenomenology in FucoPol-water isochoric supercooling systems is better understood as a mathematical discontinuity between two different physical systems with distinct characteristics: one initially predominantly anti-nucleating (Figure 8a), driven by viscosity-mediated kinetic inhibition; and another predominantly pro-nucleating (Figure 8b), driven by a conformational change that engenders the ideal microenvironment for thermodynamic Gibbs-Thomson ice modulation to manifest.

## Discussion

### Unifying hypothesis: supporting evidence

Our unified hypothesis for gel-forming, ice-binding polysaccharides agrees especially well with previous experimental and theoretical calculations. On one hand, an increased viscosity has been associated to an induction of kinetic control by establishing an anti-nucleation regime [35] and lead to stochastic narrowing [36, 37]. On the other hand, the effect of volumetric confinement due to small pore formation has been observed to result in pro-nucleation effects [38] and the observed asymmetric narrowing [39]. The Gibbs-Thomson surface binding effect expressed in polysaccharide-ice cooperativity has also led to pro-nucleation and growth modulation of cluster size [40]. However, none of these studies have reported the dual ANTI/PRO nucleation behavior, suggesting that the differentiating factor in capturing a change in system characteristics that led to a transformation of CNT system energetics was the sol-gel transition near hypothermia. Therefore, the formation of a porous gel network was discriminated as the most impactful initiator of polysaccharide cryoprotective functionality by means of phenomenological anticipation. There have been numerous studies investigating the effect of confinement on nucleation in a wide range of systems. Nanoconfined water has presented mixed literature, either agreeing with CNT formalisms [38] or showing complete nucleation suppression [41]. At the micron scale, ordered gel structures have been shown to create a spatial confinement effect which is selective towards specific crystal polymorphs, although the nucleation rate *J* can decrease or increase through a number of underlying mechanisms [39]. Researchers found that whenever the pore size was smaller than *r*\*, the growth of a polymorph of size *r*\* would be consistently hindered [42], but nucleation would be promoted when the pore dimensions were comparable to *r*\* of an emerging nucleus. Also, the pore surface area-to-volume ratio was a determinant factor in relative polymorph stability [40] and these properties largely differed from the bulk. A cross-linked polyethylene glycol diacrylate gel produced a swollen disordered polymer network which restricted molecular mobility but also decreased induction time [40]. Not only nucleated polymorphs were 3–4 times more concentrated in the gel structure than the bulk, but the growth of type II needles (small *r*\*) was promoted over type I spheroids (large *r*\*). Diao et al. [43] concluded that an increased viscosity was insufficient to explain the observations, resorting to a templating effect to explain the reduced entropic cost towards nucleation. The magnitude of this effect was remarkable, and stands equally so for our FucoPol observations, considering the disordered nature of the network when compared to 2D surfaces intentionally designed to induce specific epitaxial orientations to crystals.

### Expanded inference to other scenarios

We previously hypothesized that the fundamental driving force of cryoprotection in polysaccharides is a combination of gel-based volumetric confinement and Gibbs-Thomson specific binding. We further emphasized that no effect could disregard the other. Extending our hypothesis beyond the scope of collected data, there are certain exceptionalities to be considered. First, none of the polysaccharides we have studied [4] demonstrated a glass transition towards achieving a potential state of vitrification. Without volumetric confinement induced by a sol-gel transition, the free diffusion of water molecules through a soluble bulk would result in the growth of large crystals. It remains unknown if a polysaccharidic vitreous system could achieve similar system characteristics to a gel in terms of enabling Gibbs-Thomson binding and pro-nucleation, regardless of the proven benefits of vitrified systems in cryobiology [44]. Second, the gel environment at a macroscale is most likely not adiabatic, so nucleation promotion in a gel pocket may influence neighboring nucleation by microenvironment thermal fluctuations, or by exerting convexity changes in pore wall contact angle which disturbs ice-binding affinity. Although the observed phenomenology is expected not to change in this case, deviations from ideal conditions may exist in the balancing act of the nucleation dualism. Lastly, contrary to what was depicted in Figure 6, the walls may not be of infinitesimally small thickness *∂x* but rather form a gel-phase of higher density that confines a liquid-phase in its interstitial space. From the crystal polymorphs data obtained by Diao et al. [39], we expect an interstitial medium of higher density to be predominantly anti-nucleating, but the energetics tendency to favor growth of low *r*\* polymorphs still validates the perceived dualism observed: if a system contains compartmentalized, variable-density microenvironments, then simultaneous nucleation effects can arise, such as high-density gel-like interstitial walls with predominant anti-nucleation; and low-density, water-like pores with predominant pro-nucleation.

## Conclusion

The predominantly logical conceptualization of a cryoprotective agent is one of inducing the super-cooling of water and ice growth inhibition to avoid cryoinjury. Therefore, the cryoprotective nature of bio-based polysaccharides, particularly in our studies with FucoPol, has raised mechanistical questions over the years because it revealed that a biological post-thaw benefit to cryopreserved cells was present even under contradictory conditions of nucleation and ice growth anticipation. Moreover, the paradoxical ANTI/PRO nucleation dualism observed in FucoPol-water aqueous mixtures being cooled in a constant-volume isochoric system fueled the need for deeper understanding. In an attempt to reconcile the biological benefits of cryoprotective polysaccharides with their phenomenological effect in ice physics, we developed a three-factor mechanistical hypothesis to explain the behavior of gel-forming, ice-binding polysaccharides. These cryoprotective traits that arise from anticipating nucleation and ice growth are a combination of viscosity-mediated kinetic hampering of molecular diffusion, Gibbs-Thomson specific ice binding that induces growth modulation and size reduction, and volumetric confinement of the unfrozen water fraction in a porous gel architecture that drives the nucleating system to shift, amid cooling, from predominantly kinetic anti-nucleation to thermodynamically permissible pro-nucleation. The concerted action of these attributes explain an anticipated nucleation and ice growth, an increased stochastic determinism and the small-nuclei survivorship bias by constrained annihilation of large *r*\* growth. Classical Nucleation Theory formalisms fully support this hypothesis thermodynamically but are mathematically inconsistent with a single CNT energetic landscape, advancing that a system state change, e.g. a sol-gel transition near hypothermia, is necessary to justify the phenomenological dualism.

## CRediT Authorship

B.M.G. conceptualized study, performed experiments, data analysis, wrote manuscript. F.F., J.L., J.S. reviewed manuscript, supervised, funded. All authors have read and agreed to the published version of the manuscript.

## Data Availability Statement

The data that support the findings of this study are available from the corresponding author upon request.

