## Supplementary Information for "Elucidation of the polysaccharide cryoprotection mechanism: kinetic inhibition, thermodynamic Gibbs-Thomson modulation and volumetric confinement"

1. **Fundamentals in cryobiology**

Cryopreservation intrinsically involves a change in temperature from physiological conditions, and cellular fate during and following cold exposure is primarily determined by thermodynamic processes. Since biological processes are suppressed at cryogenic temperatures, ice nucleation and osmotic transport are the primary predictors of survival. Of all thermodynamic principles, the **First Law** and **Second Law of Thermodynamics** are most relevant to cryobiology, which concern internal energy *U* and entropy *S*, respectively. The following equation, which is a derivation of the Fundamental Relation, reflects the First Law and states that the internal energy *U* of a system is conserved:

$$dU=TdS-PdV+ \sum_{i=1}^{n} \mu_{i}dN_{i}$$

(1)

where *dU* is an infinitesimal change in internal energy, *T* is temperature, *S* is entropy, *P* is pressure, *V* is volume, *μ* is chemical potential and *N* is mole number of species *i* in a system containing *n* components. The Gibbs phase rule states that a single-component, single-state system can be completely defined by two independent intensive properties, the third being constant. The simplest relationship is the *ideal gas law*, from the known *P-V-T* relationship:

$$PV=NRT$$

(2)

where R is the *universal gas constant*. In the context of cryobiology, the *osmotic virial equation* is of greater relevance, because it relates the chemical potential *μ* of a dissolved species, such as a cryoprotectant, with the temperature, pressure and mole number in a system [1]:

$$\Pi=C_{i}+\alpha_{i}C_{i}^{2}+\beta_{i}C_{i}^{3}+\ldots$$

(3)

where Π is the total solution osmolarity, *C_i_* is the molal concentration of solute *i*, whereas α and β are the corresponding second and third osmotic virial coefficients. This equation of state can be further expanded to multi-solute systems [1]. Most thermodynamic relationships express conditions of thermodynamic equilibrium, but in cryobiology, interest is often in non-isolated systems which can interact with their surroundings. In this scenario, thermodynamic equilibrium is expressed by *free energy*, which is the system energy free to be minimized once all constraints of the interactions with the surroundings are met. For a constant-volume (isochoric) system, the thermodynamic potential is the Helmholtz free energy (Eq. 4). At constant temperature and pressure, the Gibbs free energy is defined (Eq. 5), from which the heat transfer (or enthalpy, *H*) can be derived (Eq. 6).

$$F=U-TS$$

(4)

$$G=U-TS+PV$$

(5)

$$H=U+PV$$

(6)

In cryobiology, a biological system being cryopreserved is a complex thermodynamic system made up of composite phases (ice, solution, vitrified) in both the intracellular and extracellular regions where multiple components (CPA, proteins, salts) exist; and even the phospholipid membrane can be treated as an influencing subsystem. One last important relationship is that of interfacial tension *γ* between two surfaces, as a change of internal energy *U* by interacting surface area *A*:

$$\gamma= \left( \frac{\partial U}{\partial A} \right)_{S, N_{1},N_{2},\ldots,N_{n}}$$

(7)

Interfacial thermodynamics are highly relevant in cryoprotectant-ice and cryoprotectant-membrane interactions and explain the widely documented adsorption-inhibition mechanism in antifreeze proteins, with TH as its thermodynamic manifestation; and the IRI mechanism.

1.1. **Nucleation**

A pure liquid with a melting temperature T*_m_* being cooled is defined by a Gibbs free energy for both its solid (*S*) and liquid (*L*) states, and the following relationship applies:

- At $\text{T}\text{ > }\text{T}_{\text{m}}$ the liquid state is thermodynamically favored ($\text{G}_{\text{L}}\text{ < }\text{G}_{\text{S}}$).
- At $\text{T}\text{ < }\text{T}_{\text{m}}$ the solid state is thermodynamically favored ($\text{G}_{\text{L}}\text{ > }\text{G}_{\text{S}}$).

The liquid-solid phase transition is a first-order exothermic phase transition, because $\text{∂G/∂T}$ is discontinuous at $\text{T}_{\text{m}}$ and $\text{∆H<0}$, respectively. It then follows that at:

$$T= T_{m} \underset{\to}{} \Delta G =0 \underset{\to}{} \Delta S={\Delta H}/{T_{m}<0}$$

A negative change in entropy reflects the reduction in the rate of molecular translational and rotational operations from liquid to solid. When assuming that $\text{∆H}$ is independent of $\text{T}$, a new equilibrium freezing point of kinetic nature can be expressed by $\text{∆G}$, as a function of a thermal hysteresis $\text{TH}$ centered around T*_m_*:

$$\Delta G(T)=TH\frac{\Delta H}{T_{m}}$$

(8)

$$TH=T-T_{m}$$

(9)

The concept of thermal hysteresis indicates that, thermodynamically, water should freeze at
T = T*_m_* but it relies on kinetic processes such as molecular diffusion and clustering to produce crystals, hence creating a temperature lag. Logically, crystallization should occur whenever
T < T*_m_*, but it is a non-spontaneous process ($\text{Δ}\text{G>0}$) due to a decrease in entropy. Experimentally, water does not freeze at 0 °C, and very pure water has shown to be kinetically stable down to temperatures of *ca.* –40 °C [2]. Thus, during the cooling window, a sub-T*_m_* liquid is said to be in a metastable supercooled state and requires a driving force to kickstart the spontaneous process. That driving force is called nucleation. The nucleation process initiates (Figure 5 on the main paper) when a small agglomerate of water molecules (an embryo, or cluster) begins to form in the liquid as a result of minimized molecular collisions, as attractive interactions begin dominating over repulsive forces. This cluster is unstable and may continuously grow (as free water molecules migrate to it) or re-dissolve into the environment. The embryonic (i.e. unstable cluster) stage of nucleation is then an energetic balance between the free energy available from the driving force, and the energy consumed in forming a new interface, and occurs for nuclei radii 0 < *r* < *r** (critical radius). For nuclei radii *r** < *r* < *r_eq_* (equilibrium radius), stable clusters continuously grow without redissolution, albeit under non-spontaneous thermodynamic potential. Once the rate of change of free energy becomes negative, then an embryo can grow [3]. Thus, a gain-cost function creates an energetic landscape from which all properties of nucleation and crystallization can be derived (Figure 5 on the main paper). The crucial point is to understand it as a balance between the free energy available from the driving force (volumetric free energy, VFE), and the energy consumed in forming new interface (interfacial energy, IE).

**1.2.** **Boundary conditions for nucleation**

The growth of a cluster $\text{A}_{\text{n}}$ at T < T*_m_* does not spontaneously result in stable nuclei. In the initial phases, a growth-decay stage governed by thermal fluctuations generates local aggregations of water molecules but does not constitute a two-phase system. The transition from unstable clusters (below *r**) to stable nuclei (above *r**) is governed by a potential energy landscape and arises at an inflection point in free energy. The nucleation free energy is obtained by a chemical potential difference in the parent and equilibrium phases for all atoms in the cluster:

$$\Delta G=\sum_{i} y_{i}^{e}\left( \mu_{i}^{e}-\mu_{i}^{0} \right)$$

(10)

where $\text{y}_{\text{i}}^{\text{e}}$ is the atomic fraction of atom *i* in the nucleating equilibrium phase, while $\text{μ}_{\text{i}}^{\text{e}}$ and $\text{μ}_{\text{i}}^{\text{0}}$ are the chemical potentials in the nucleating and parent phases, respectively. When the parent phase is metastable, the chemical potential is higher than at equilibrium, becoming the driving force of nucleation as $\text{∆G}$ becomes negative.

**1.3.** **Homogenous nucleation**

Homogenous nucleation occurs when pure water nucleates from water molecules alone. This implies that a spherical cluster is the only driving force for nucleation to occur. The nucleation activation energy barrier $\text{∆G}_{\text{n}}$ is an energetic balance between opposing contributions. When a cluster of a new phase forms, the system decreases its free energy. This is the driving force for nucleation and is directly proportional to the volume of the cluster:

$$\text{VFE} = -\frac{4}{3}\pi r^{3}{\Delta G}_{v}$$

(11)

A two-phase system implies the creation of an interface between the parent phase and the new cluster. This has an energetic cost, proportional to the surface area of the cluster:

$$\text{IE} = 4\pi r^{2}\gamma_{SL}$$

(12)

The nucleation activation energy barrier $\text{∆G}_{\text{n}}$ can then be expressed as a combination of energetic gain (VFE, or $\text{∆G}_{\text{v}}$) and cost (IE, or $\text{γ}_{\text{SL}}$) functions, as follows:

$$\text{∆G}_{\text{n}}\text{ = VFE + IE} =-\frac{4}{3}\pi r^{3}{\Delta G}_{v}+4\pi r^{2}\gamma_{SL}$$

(13)

where $\text{∆G}_{\text{v}}$ is the volumetric free energy and $\text{γ}_{\text{SL}}$ is the solid-liquid interfacial energy. The preceding coefficients of both energy parameters correspond to the volume and surface area of a spherical cluster, respectively, produced by homogenous nucleation. Eq. 13 can also be expressed as a function of number of atoms *n* in a cluster:

$$\text{∆G}_{\text{n}}= n{\Delta G}_{v}+n^{2/3}A\sigma$$

(14)

where, $\text{σ}$ is the interfacial tension and $\text{A}$ is a geometric factor (Figure SI.1).

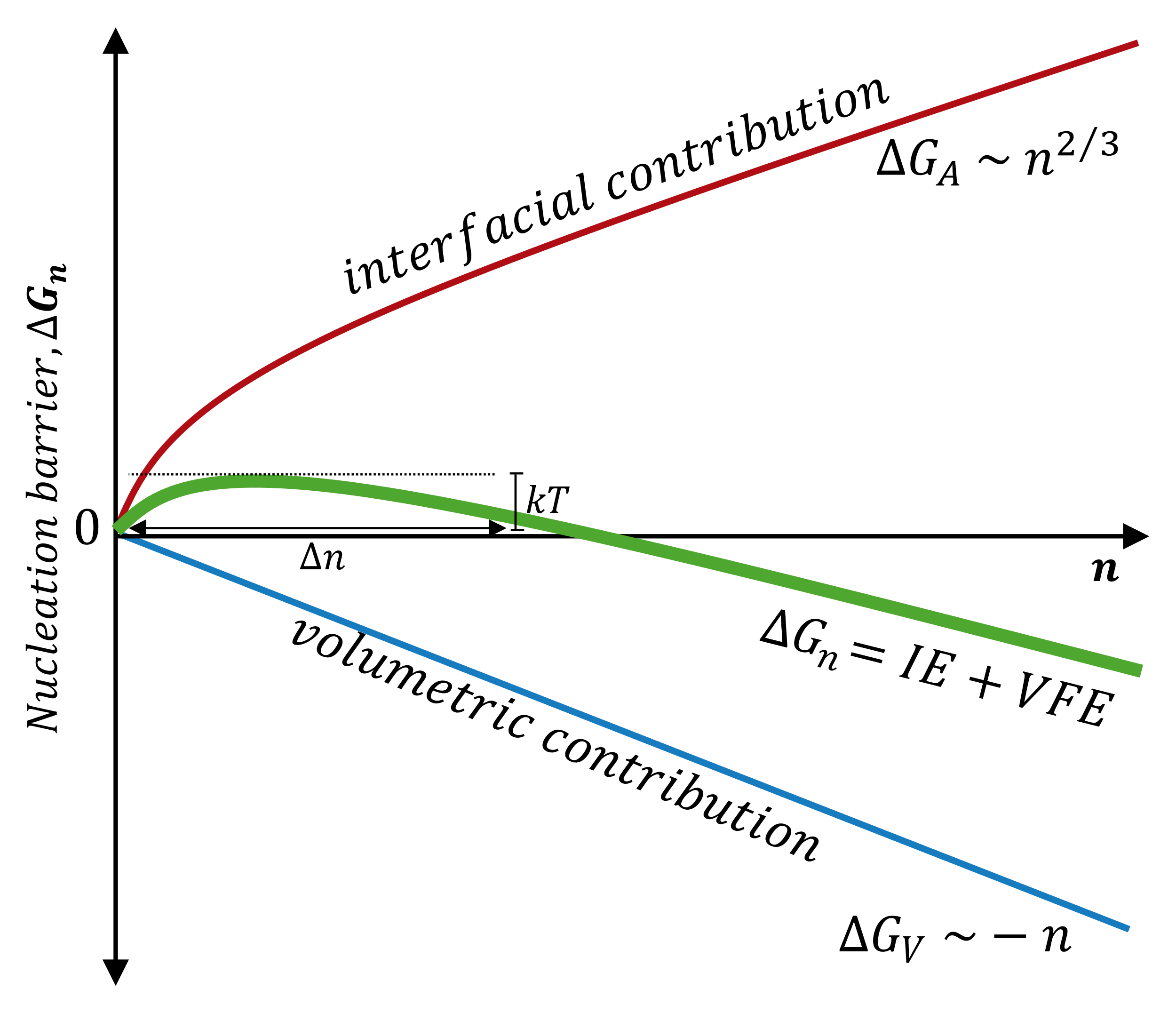

Figure SI.1 – Influence of system scaling *n* in the energetic components driving nucleation.

**1.4. Heterogenous nucleation**

Heterogenous nucleation occurs at preferential binding sites of a dissolved molecule, in phase boundaries, container surfaces or impurities and occurs much more often than homogeneous nucleation. The formalisms are essentially the same between homogenous and heterogenous nucleation, but the latter considers a contact angle component (Figure SI.2).

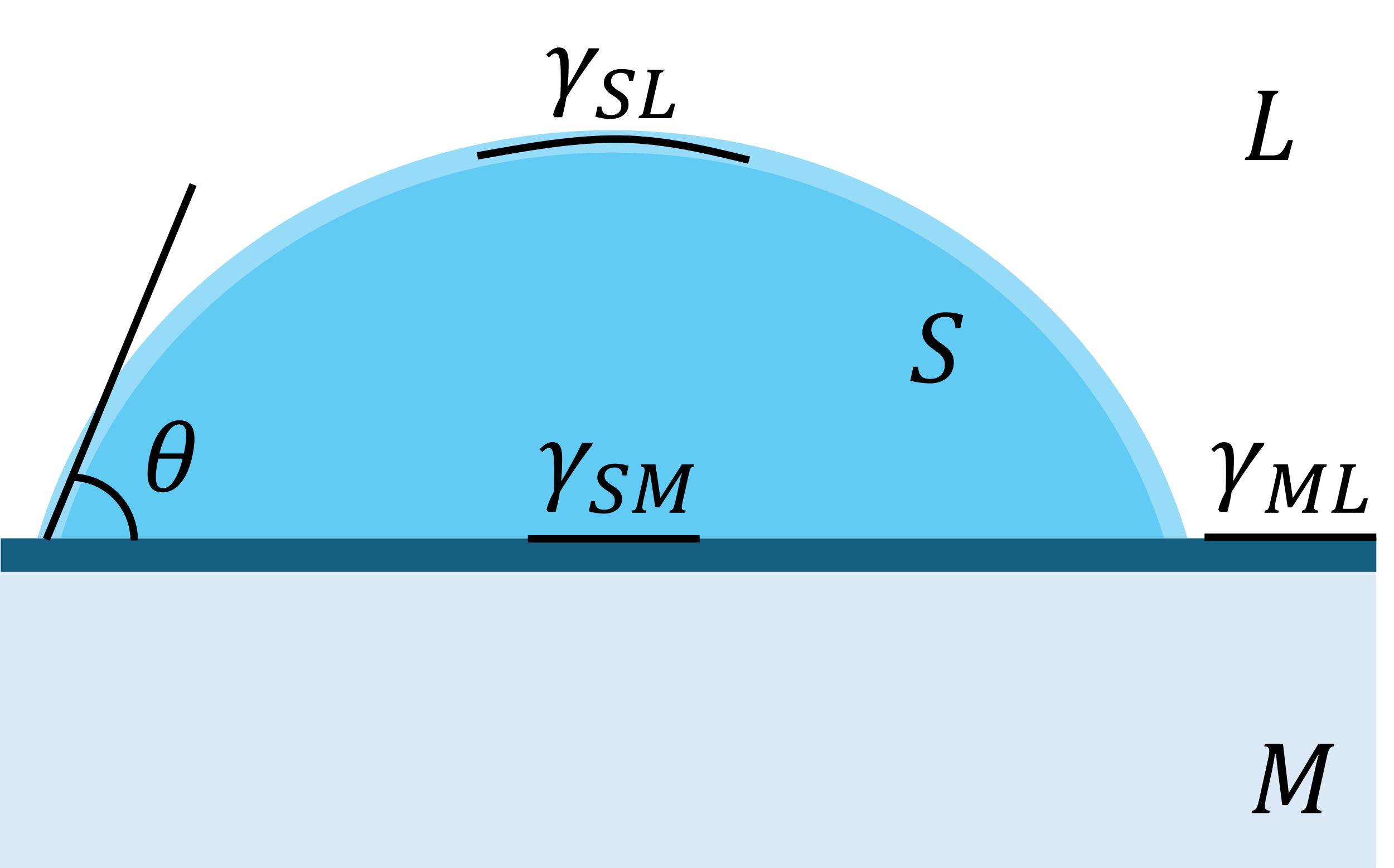

Figure SI.2 – Visual representation of energetic components *γ* acting on a liquid droplet of contact angle *θ*.

A cluster *S* in contact with the specific surface area of a molecule *M* in a liquid system *L* creates a contact angle *θ* originating from the intersection of tangential surface tension vectors between each pair of phases $\text{γ}_{\text{SM}}$, $\text{γ}_{\text{SL}}$ and $\text{γ}_{\text{ML}}$ and is defined by the Young equilibrium equation:

$$\cos\theta=\frac{\left( \gamma_{ML}-\gamma_{SM} \right)}{\gamma_{SL}}$$

(15)

Without the need to generate a spherical cluster *ex nihilo*, these surfaces promote nucleation due to wetting, in which small contact angles ($\text{θ < π}$) convert to a uniform distribution of the *n* atoms in a cluster along the specific surface area, facilitating nucleation by reducing the interfacial energy component (Figure 8 in the main paper, Eq. 12). Hence, $\text{ΔG}_{\text{n}}^{\text{het}}\text{<}\text{ ΔG}_{\text{n}}^{\text{hom}}$. A contact angle $\text{θ=π}$ arises when the interface free energies between the liquid and the cluster do not allow for wetting of the surface *M* to proceed, as an unwet cluster costs less energy. However, if $\text{γ}_{\text{ML}}$ – $\text{γ}_{\text{SM}}$ is greater than $\text{γ}_{\text{SL}}$ then wetting can occur. Heterogenous nucleation is then an extension of Eq. 13, by implementing a trigonometric component $\text{f}\left( \text{θ} \right)$ into the original formalism:

$${\Delta G}_{n}^{het}={\Delta G}_{n}^{hom}\cdot f\left( \theta\right)= \left[ -\frac{4}{3}\pi r^{3}{\Delta G}_{v}+4\pi r^{2}\gamma_{SL} \right]\left[ \frac{2-3\cos\theta+\cos^{3} \theta}{4} \right]$$

(16)

**1.5. Critical conditions for stable nucleation**

An unstable cluster will only become energetically stable when a critical radius *r** is achieved, past which point it can grow without decaying. From the energetic landscape of Figure 5 (main paper):

- when $\text{r < }\text{r}^{\text{*}}$, an increase in *r* leads to an increase in $\text{∆G}_{\text{n}}$ creating unstable embryos that cyclically cluster and redissolve proportionately to thermal fluctuations.
- when $\text{r ≥ }\text{r}^{\text{*}}$, an increase in *r* leads to a decrease in $\text{∆G}_{\text{n}}$ creating stable nuclei that can grow from size *r** to *r_eq_* following a Gaussian distribution.

Likewise, the critical radius, *r** is associated to a critical free energy for nucleation, $\text{Δ}\text{G}_{\text{n}}^{\text{*}}$. When the first derivative $\frac{\text{d}\left( \text{Δ}\text{G} \right)}{\text{dr}}\text{=0}$, the boundary conditions for stable nucleation to occur and progress into crystallization can be associated to thermodynamic parameters, such as melting temperature, thermal hysteresis and solid-liquid interfacial energy:

$$r^{*}= \frac{2\gamma_{SL}}{{\Delta G}_{v}}=\frac{1}{\Delta T}\left( \frac{2\gamma_{SL}T_{m}}{L_{V}} \right)$$

(17)

$$\Delta G_{n}^{*}= \frac{16\pi\gamma_{SL}^{3}}{3{\Delta G}_{V}^{2}}=\frac{1}{\left( \Delta T \right)^{2}}\left( \frac{16\pi\gamma_{SL}^{3}T_{m}^{2}}{3L_{V}^{2}} \right) = \frac{4}{27}\frac{\left( A\gamma_{SL} \right)^{3}}{\left( {\Delta G}^{nuc} \right)^{2}}$$

(18)

These fundamental relations constitute the widely accepted **Classical Nucleation Theory** (CNT). Its landmark assumption is that nucleation is a stochastic process, *i.e.* a probabilistic event for which an accurate temperature interval can be pinpointed, but a given T*_n_* cannot be predetermined. Thus, a cluster of radius near *r** is not guaranteed to become stable when $\text{Δ}\text{G}_{\text{n}}^{\text{*}}$ is achieved. The probabilistic aspect of nucleation is mediated by local temperature fluctuations arising from the frequency of molecular collision and is represented by the Zeldovich factor, *Z*:

$$Z=\frac{3\left( {\Delta G}_{n} \right)^{2}}{4\sqrt{\pi kT}\left( A\sigma\right)^{3/2}}$$

(19)

Near $\text{Δ}\text{G}_{\text{n}}^{\text{*}}$ the energy profile has an associated uncertainty *Z* arising from thermal fluctuations *kT*, which reflects the flatness of the curve. For two systems having the same free energy barriers, the system with the flatter free energy landscape near the barrier has more diffusive nucleation dynamics and a lower nucleation rate.

**1.6. Nucleation rate**

If crystallization is a kinetic problem, one can assume that it is defined by rates of change *k*, whereas freezing should be faster at –40 °C than at –0.1 °C. As T decreases, the loss of molecular mobility implies that water molecules will start aggregating into small clusters. By the law of mass action, one can then define a small molecular cluster *A_n_* with *n* molecules of species *A* that grows (*A_n_*_+1_) or decays according to association or dissociation rate constants *k_a_* and *k_d_* by addition to, or loss of one molecule from, *A*:

$$A_{n}+A_{1}\begin{matrix} k_{a} \\ \rightleftharpoons\\ k_{d} \end{matrix}A_{n+1}$$

From a system state population perspective, nucleation generates stable cluster sizes that follow a Boltzmann distribution. The computation of population densities is important to discuss temperature-domain and time-domain metrics (Figure 6 in the main paper), such as the nucleation temperature T*_n_* and nucleation rate *J*, respectively. A population of clusters near the critical regime is given by:

$$n_{r}= n_{0}\cdot\exp\left( -\frac{{\Delta G}_{r}}{kT} \right)$$

(20)

where *k* is the Boltzmann factor, $\text{n}_{\text{0}}$ is the number of atoms in the system and $\text{ΔG}_{\text{r}}$ is the free energy associated with the cluster of size *r*. At the critical radius *r**, the population density of clusters of radius *r** (*C**) is:

$$C^{*}= C_{0}\cdot\exp\left( -\frac{{\Delta G}_{n}^{*}}{kT} \right) \text{clusters}\text{/}\text{m}^{\text{3}}$$

(21)

where $\text{C}_{\text{0}}$ is the atomic fraction accessible to the cluster (for precipitation in the solid state, $\text{C}_{\text{0}}\text{=1}$). The addition of one more atom to each of these clusters would convert them to stable nuclei at a frequency *f*_0_. When the nucleation barrier $\text{Δ}\text{G}_{\text{n}}^{\text{*}}$ is greater than *kT* the system can overcome its metastable character and allow a critical-size nucleus to grow at a steady-state, given by the nucleation rate, *J*:

$$J_{hom}= f_{0}\cdot C_{0}\cdot\exp\left( -\frac{{\Delta G}_{hom}^{*}}{kT} \right) \text{nuclei}\text{/}\text{m}^{\text{3}}$$

(22)

$$J_{hom}= f_{0}\cdot C_{0}\cdot\exp-\left( \frac{A}{kT}\frac{1}{\left( \mathrm{TH} \right)^{2}} \right) \text{nuclei}\text{/}\text{m}^{\text{3}}$$

(23)

$$A= \frac{16\pi\gamma_{SL}^{3}T_{m}^{2}}{3L_{V}^{2}}$$

(24)

By analogy, the heterogenous nucleation rate is then:

$$J_{het}= f_{1}\cdot C_{1}\cdot\exp\left( -\frac{{\Delta G}_{het}^{*}}{kT} \right) \text{nuclei}\text{/}\text{m}^{\text{3}}$$

(25)

**1.7. Crystallization**

Briefly, the initial conditions of nucleation pre-determine the onset and extent of crystallization. Following the energetic profile of Figure 5 (main paper), and recalling that crystallization becomes a spontaneous process whenever $\text{∆G<0}$, all stable nuclei produced within predictable size distributions by the values of *r*, *r** and *r_eq_* are allowed to grow to system size. The temperature at which perceived crystallization occurs, *i.e.* the macroscopic freezing point, is therefore a consequence of the preceding nucleation mechanisms. At this stage, the ice-free supercooled liquid state transitions into a solid state until thermodynamic equilibrium is achieved, which corresponds to a total conversion from liquid to solid in one-phase isobaric systems, or a partial conversion in two-phase isochoric systems.

2. **FucoPol, the model cryoprotectant polysaccharide**

FucoPol is a high-molecular-weight (1.7−5.8×10^6^ Da) fucose-containing, polyanionic, cryoprotective extracellular polysaccharide (EPS) produced by the Gram-negative bacterium *Enterobacter* A47 (DSM 23139) [4]. FucoPol has a fucose, galactose, glucose, and glucuronic acid hexamer motif (2.0:1.9:0.9:0.5 M ratio), a main chain composed of a →4)-α-L-Fuc*p*-(1→4)-α-L-Fuc*p*-(1→3)-β-D-Glc*p*(1→ trimer repeating unit, and a trimer branch α-D–4,6-pyruvyl-Gal*p*-(1→4)-β-D-GlcA*p*-(1→3)-α-D-Gal*p*(1→ in the C_3_ of the first fucose [5]. It also contains 13−14 wt.% pyruvyl, 3−5 wt.% acetyl, and 2−3 wt.% succinyl in its composition [4]. The presence of glucuronic acid as well as the acyl substituents pyruvyl and succinyl, confer a polyanionic character to the biopolymer [4]. It has shown strong non-colligative ice growth disruption and re-shaping [5], nucleation promotion and stochastic narrowing [6], versatile use in cryoprotective formulas of variable composition and different cell lines [7], antioxidant [8] and photoprotective effects that counteract the reactive oxygen species-mediated, cryopreservation-induced MPTP opening that leads to delayed post-thaw cell death [9]. All these properties, associated with a high molecular weight, high viscosity and shear-thinning behavior [10] that facilitates nutrient diffusion and promotes cellular attachment by natural design [11], conceive a molecule capable of maximizing the post-thaw viability of cryopreserved biologicals.

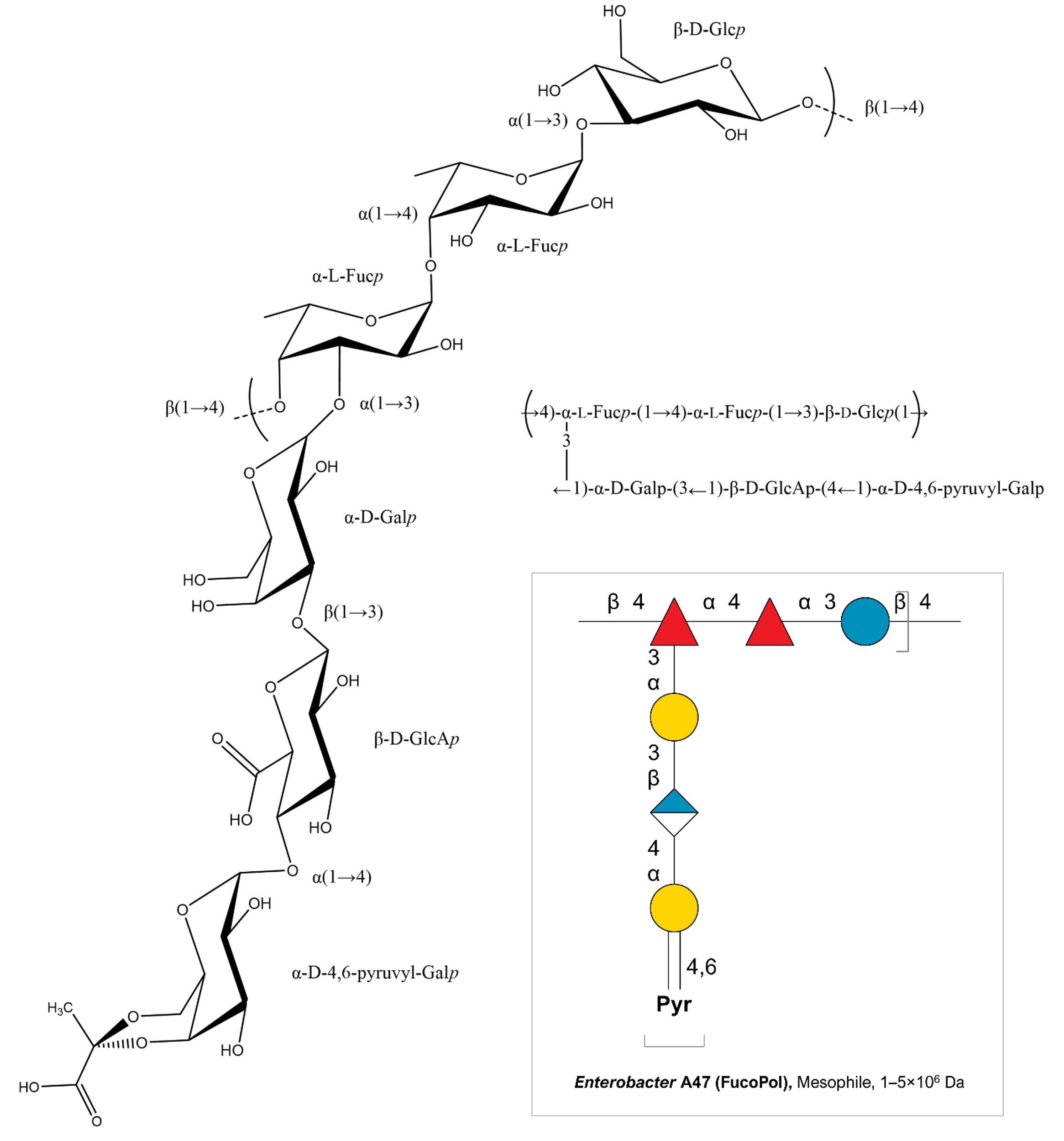

Figure SI.3 – NMR-elucidated structure of the bacterial polysaccharide FucoPol. It has a fucose, galactose, glucose, and glucuronic acid hexamer motif (2.0:1.9:0.9:0.5 M ratio), a main chain composed of a →4)-α-L-Fucp-(1→4)-α-L-Fucp-(1→3)-β-D-Glcp(1→ trimer repeating unit, and a trimer branch α-D–4,6-pyruvyl-Galp-(1→4)-β-D-GlcAp-(1→3)-α-D-Galp(1→ in the C_3_ of the first fucose. FucoPol also contains 13−14 wt.% pyruvyl, 3−5 wt.% acetyl, and 2−3 wt.% succinyl in its composition. The structure presented is a deacetylated, desuccinylated form of the biopolymer.
